# Single-cell phylodynamic inference of tissue development and tumor evolution with scPhyloX

**DOI:** 10.1101/2024.05.15.594328

**Authors:** Kun Wang, Zhaolian Lu, Zeqi Yao, Xionglei He, Zheng Hu, Da Zhou

## Abstract

Phylodynamics inference (PI) is a powerful approach for quantifying population dynamics and evolutionary trajectories of natural species based on phylogenetic trees. The emergence of single-cell lineage tracing technologies now enables the reconstruction of phylogenetic trees for thousands of individual cells within a multicellular organism, opening avenues for employing PI methodologies at the cellular level. However, the intricate process of cell differentiation poses challenges for directly applying current PI frameworks in somatic tissues. Here, we introduce a novel computational approach called single-cell phylodynamic explorer (scPhyloX), designed to model structured cell populations in various cell states, by leveraging single-cell phylogenetic trees to infer dynamics of tissue development and tumor evolution. Our comprehensive simulations demonstrate the high accuracy of scPhyloX across various biological scenarios. Application of scPhyloX to three real datasets of single-cell lineage tracing unveils novel insights into somatic dynamics, such as the overshoot of cycling stem cell populations in fly organ development, clonal expansion of multipotent progenitors of hematopoiesis during human aging, and pronounced subclonal selection in early colorectal tumorigenesis. Thus, scPhyloX is an innovative computational method for investigating the development and evolution of somatic tissues.

## Introduction

The development and maintenance of multicellular tissues depend on the specific functions and dynamics of various cell types that form hierarchical structures^1-4^. Understanding the growing dynamics of each cell type, the cell-fate decision of division and differentiation that contribute to normal tissue renewal, the somatic evolution during cancer initiation are key biological questions.

Single-cell lineage tracing has been a powerful approach for studying developmental dynamics. Somatic mutations reveal the ancestral relationships among individual cells and the dynamic processes of reproduction and differentiation. We can leverage endogenous or artificially induced somatic mutations to reconstruct cell phylogenies and infer developmental dynamics^5-9^. The recent advancements in utilizing CRISPR-Cas9 editing to track cell lineages offer a chance to reconstruct the cell lineage tree in high throughput and at whole-organism or whole-organ level^10-13^. Ideally, decoding developmental process might require complete lineage data including both all current and ancestral cells. However, in reality a developmental tree only include cells from a snapshot sample. Therefore, a comprehensive understanding of developmental dynamics with lineage tracing data sampled at a single time point requires sophisticated computational and predicative methods.

In fact, the topology and statistical properties (e.g. branch lengths) of a phylogenetic tree encodes the population dynamics in an evolutionary process^14^. This approach is known as phylodynamics inference (PI) ^15^, which aims to infer the dynamics of a population via phylogenetic trees. PI was originally developed for epidemiological studies to infer virus transmission dynamics from genomic data^16-19^. Subsequently, this method has been extended to other biological systems, such as tissue development and tumor growth^9, 20-22^. In previous studies, fixed-parameter models were commonly employed. For example, TreeTime^23^, a python package, is often used to estimate molecular clock phylogenies and population size histories using maximum likelihood approach. BEAST^24^ used fixed-parameter models like the chi-square Markov model for analysis, with limited exploration of models accounting for parameter variation over time. Werner et al.^21^ estimated the mutation rate and branch division probability using bulk sequencing data, along with patient specific evolutionary parameters in human cancers. TiDeTree^25^ enables joint inference of the time-scaled trees and cell population dynamics. These studies often assumed constant dynamic parameters or treated all cells as identical cell types with uniform dynamics, which may not accurately reflect practical scenarios where a tissue typically consists of cells in different states or types.

Here, we introduced a novel PI model, <u>s</u>ingle-<u>c</u>ell <u>phylo</u>dynamics e<u>x</u>plorer (scPhyloX), characterized by structured cell population and time-varying parameters. We also developed parameter estimation methods to infer the dynamics of stem and differentiated cells during tissue development. By analyzing the lineage tracing datasets of embryonic development from 9 organs in 2 fruit flies, hematopoietic stem and progenitor cells from 8 human donors as well as neoplastic cells from 8 mouse colorectal tumors, scPhyloX was able to identify interesting patterns of tissue development and somatic evolution. For instance, cycling stem cells show a generally overshooting population size in fruit fly organ development, which might be a mechanism of rapid embryo growth that prevents cell death. In human hematopoiesis, the ratio of progenitor to stem cells increase along human ageing. Finally, scPhyloX inference with large cell phylogenetic trees revealed strong subclonal selection during early tumorigenesis of colon cancer.

## Results

### The framework of scPhyloX for inferring developmental dynamics

All multicellular organisms are derived from a single zygote through division and differentiation. This developmental process naturally results in a cell phylogenetic tree, in which zygote lies at the root and phylogenetic branches represent cell divisions. A complete and accurate cell phylogenetic tree allows precise assessment of cell divisions and differentiation by recording the duration of cell division and the types of cells produced (**Fig. 1a**). However, reconstructing cell phylogenetic trees is typically done retrospectively using somatic mutations. The unobservable nature of cell division and other factors such as sampling, cell death, etc. lead to incomplete cell lineage data. The cells we observe are the leaves of the phylogenetic tree, and all the internal nodes are pseudo-progenitors inferred in the phylogenetic tree reconstruction. Despite the incomplete nature of the reconstructed cell lineage and the loss of information regarding internal node states, information from the observed cells can still offer insights into overall tissue development dynamics. The rate of somatic mutations and cell death together determine the distribution of branch lengths from the leaf nodes to their immediate internal notes, defined here as the leaf-progenitor (LP) distance (*ξ*). The rates of cell division, differentiation and death determine the distribution of the branch lengths from the leaf nodes to root, defined here as the leaf-root (LR) distance (*η*) (**Fig. 1b**). By deriving the distributions of *ξ* and *η*, we can estimate phylodynamics parameters from the phylogenetic tree.

**Fig. 1.**
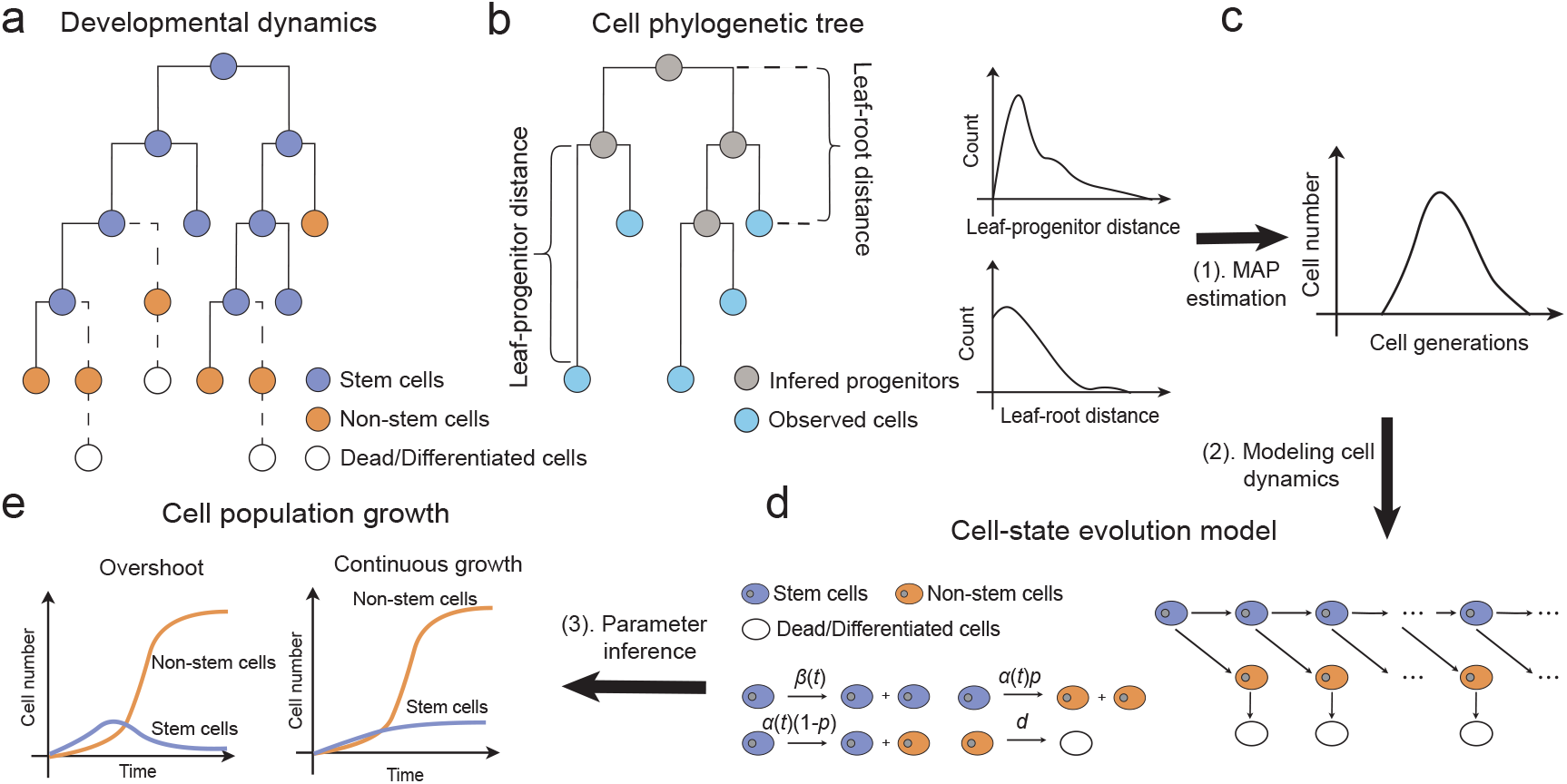
Schematic of scPhyloX for inferring developmental dynamics from single-cell lineage tracing data. (**a**), Developmental dynamics of a cell hierarchy including stem cells and non-stem cells. A stem cell (purple) can either self-replicate or differentiate into non-stem cells (orange). Non-stem cells either remain non-dividing state or die leading to lose their lineages at a particular probability (dashed lines). (**b**), A cell phylogenetic tree reconstructs the cell lineage relationship for a sample of cells. Blue circles indicate the observed cells (leaves) while grey circles indicate the unobserved progenitors (internal nodes). Here, we modeled the distributions for distance from an observed cell to its most recent progenitor (denoted as leaf-progenitor distance) and the distance to the root cell (denoted as leaf-root distance), respectively. (**c**), MAP (maximum-a-posteriori) estimation of cell numbers per generation from the distributions of leaf-progenitor distances and root distances. (**d**), Cell-state evolution model. In each generation, stem cells (purple) can either divide symmetrically generating two stem cells or two non-stem cells (orange), or asymmetrically generating one stem cell and one non-stem cell. (**e**), Line graph showing cell population dynamics under two different growth modes. In overshoot mode (left), the number of stem cells during growth can exceed the final static number at tissue homeostasis. In continuous growth mode (right), the number of stem cells is increasing all the time until reaching the final static number at homeostasis. In both modes, the growth of non-stem cells is monotonic with a plateau (S-shape).

Under the assumption of infinite site model^21, 26^, we can calculate the distribution of LP distance in phylogenetic tree (**Methods**) as,

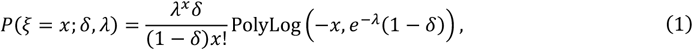

where *λ* is mutation rate, *δ* is the probability of lineage loss due to death or differentiation, PolyLog(*n, z*) is polylogarithm function *Li*_*n*_(*z*)^27^.

To measure the LR distance, we first need to estimate the number of cells in each generation (***x*** = (*x*_1_, ⋯, *x*_*n*_)^T^). Utilizing the mutation rate *λ* estimated by equation (1), the LR distance for i-th generation cells *η*_*i*_ follows a Poisson distribution,

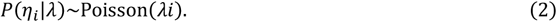

Then the number of cells in i-th generation *x*_*i*_ can be estimated using maximum a posteriori (MAP) estimation (**Fig. 1c, Methods**).

To relate the LP and LR distances to developmental dynamics, we present a dynamic model that describes the state transition of cells within tissue development, accounting for the distinct functions of stem and non-stem cells. The fate of stem cells is determined by the probabilities of self-renewal, *β*(*t*), leading to the generation of two offspring stem cells, or differentiation, 1 − *β*(*t*), which results in the cessation of the cell division. We consider two modes of division during stem cell differentiation: symmetric division, with probability *p*, yielding two non-stem cells, or asymmetric division with probability 1 − *p*, producing one stem and one non-stem cell. Non-stem cells are subject to a death rate of *d*. We denote the population sizes of stem and non-stem cells as *SC* and *NC*, respectively, and employ dynamic equations to describe the temporal evolution of these populations, providing insights into the phylodynamics of cell count as influenced by the parameters,

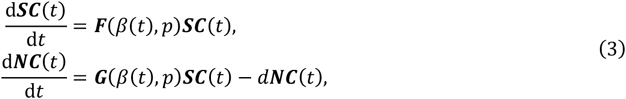

where ***S****C*(*t*) = *SC*_0_(*t*), ⋯, *SC*_*n*_(*t*))^*T*^, ***N****C*(*t*) = (*NC*_0_(*t*), ⋯, *NC*_*n*_(*t*))^T^, *SC*_*i*_ (*t*) and *NC*_*i*_ (*t*) are the numbers of stem cells and non-stem cells in the i-th generation at time *t*, ***F*** and ***G*** are coefficient matrices of the equations (**Fig. 1d**). By solving **Eq. (3)**, the total number of cells in the i-th generation at time *t* can be calculated as *x*_*i*_ (*t*) = *SC*_*i*_ (*t*) + *NC*_*i*_ (*t*). Combined with **Eq. (2)**, we can estimate the dynamic parameters of tissue development and thereby infer the dynamic history of tissue development. (**Methods**)

Upon analyzing the parameters of the dynamic equations, we observed that the growth of stem cells in tissues follows two different models, determined by a parameter designated as the developmental dynamics index (DDI) *b*^*^. When DDI *b*^*^ < 1, the quantity of stem cells initially rises and subsequently falls, a pattern termed overshooting growth. Conversely, when DDI *b*^*^ ≥ 1, the population of stem cells exhibits sustained growth, a behavior described as continuous growth (**Fig. 1e, Methods**).

### ScPhyloX recovers cell population dynamics in simulations and benchmarked cell line

Using our previously published simulation framework^28^, we performed stochastic simulations of somatic mutation accumulation in a growing cell population. We used the Gillespie algorithm^29^ to simulate the cell birth, differentiation and death events. At each division, cells acquired a number of new mutations following a Poisson distribution with given expectation *λ*. The simulation ended when the simulation time reached the preset value (**Methods**). We then randomly sampled 500 single cells to reconstruct the phylogeny using the accumulated mutations in individual cells and estimate the population dynamics parameters with scPhyloX. We modeled two scenarios of tissue dynamics with distinct growing trajectories of stem and non-stem cells, the overshoot model and the continuous growth model (**Fig. 2a-d**), respectively. In overshoot model, stem cells first proliferate through division, expanding their numbers to a high level, and then gradually differentiate into non-stem cells, showing a pattern in which the number of stem cells first increases and then decreases. The overshoot model was found in previous studies on development of intestinal stem cells and development of neocortical neurons^4, 30^. While in continuous growth model, stem cells simultaneously divide and differentiate, with the division rate marginally exceeding the differentiation rate. Eventually, the rates of division and differentiation stabilize, leading to a steady increase and maintenance of the stem cell population. Continuous growth model was found in previous study to be able to maintain stem cell populations, such as crypt stem cell development using this strategy ^4^.

**Fig. 2.**
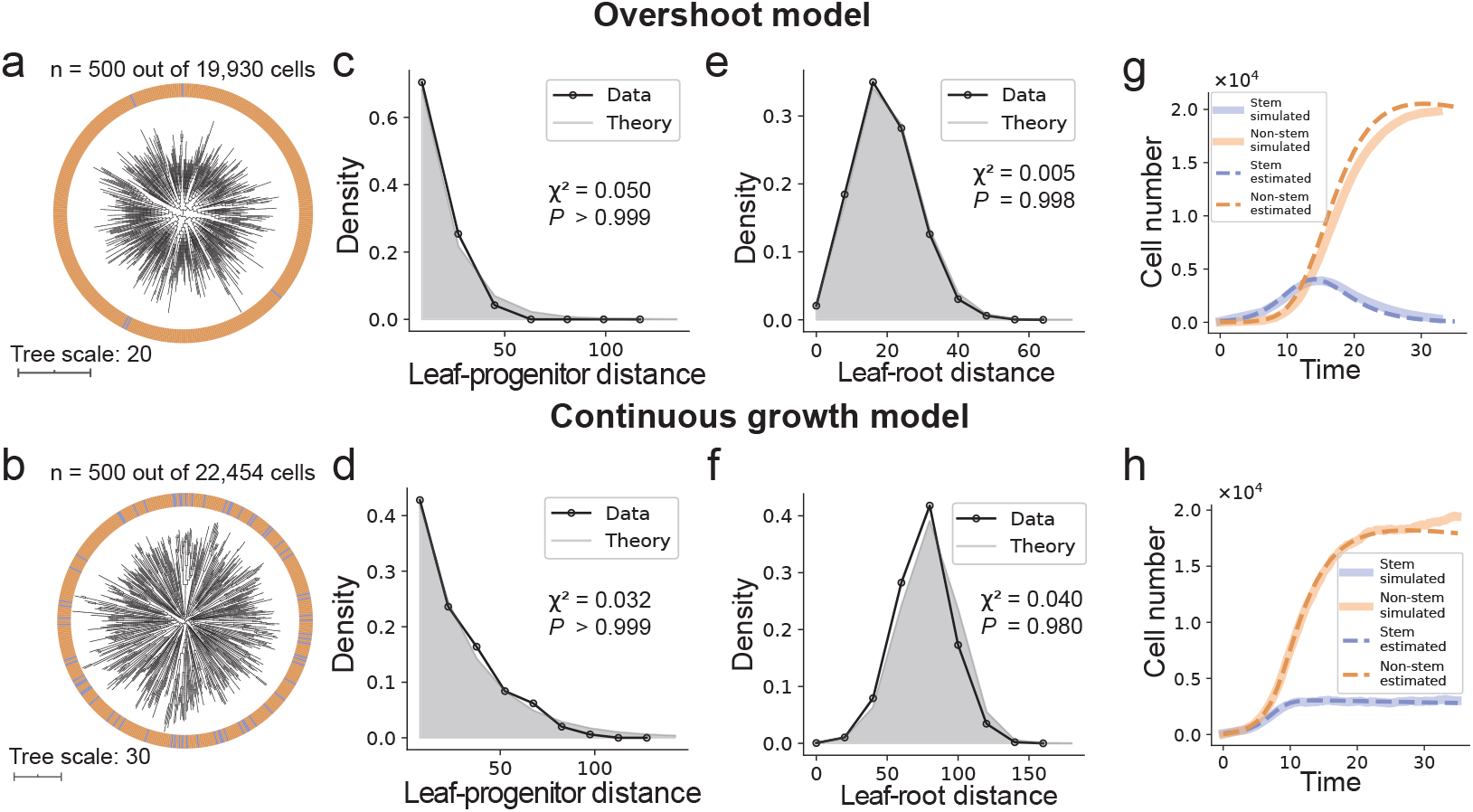
scPhyloX accurately recovers developmental dynamics in simulations. (**a-b**), Single-cell phylogenetic trees generated under simulated overshoot model (**a**) and continuous growth model (**b**) of tissue development, respectively. (**c-d**), The model fitting of the distribution of leaf-progenitor distances in the two models. (**e-f**), The model fitting of the distribution of leaf-root distances in the two models. Pearson’s chi-square tests are shown here. (**g-h**), Inferred cell population growth (dashed lines) matches simulations (solid lines) in both models. Stem and non-stem cells are purple and orange, respectively.

To optimize the estimations of numerous parameters in scPhyloX, we devised a combined optimization method for parameter inference. We applied the differential evolution (DE)^31, 32^ algorithm to stochastically search for the global optimum, and then used the differential evolution Markov Chain Monte Carlo (DE-MCMC)^33, 34^ method to obtain the parameter distributions (**Methods**). This method allowed us to accurately estimate parameters and give interval estimation.

We found that the inference results truly reflect the dynamic pattern of overshoot and continuous growth of stem cells (**Fig. 2c-d**). The ground truth values of each simulated parameters fall within the middle range of the inferenced distribution (convergence diagnostic^35^ 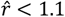 in most parameters) (**Fig.2e-h, Supplementary Fig1, 2, Supplementary Table 1)**.

To validate the performance of scPhyloX in real data, we first applied the method to a high-resolution single-cell phylogeny mapped by DNA Ticker Tape technique in *in vitro* culture of HEK293T cell line^36^. The HEK293T cells in the tree were sampled from a single-cell-derived clone. Because this immortalized cell line is considered non-differentiated, it mainly consists of cycling “stem-like” cells. In fact, the LR distance in the phylogenetic tree fitted a normal distribution well, indicating a highly similar proliferative rate of the cells. Therefore, exponential growth with only cycling stem cells would be expected. Indeed, although we did not deliberately set the growth pattern of exponential growth for the model, scPhyloX correctly inferred the exponential growth pattern of HEK293T cells and inferred that almost all cells were at the cycling state (**Supplementary Fig 3**).

### The overshooting growth of stem cells in fly organ development

We next applied scPhyloX to single-cell lineage tracing data in two fruit fly embryos, recorded using SMALT (Substitution Mutation-Aided Lineage Tracing), which is a high-resolution phylogenetic mapping method enabled by AID (activation deamination)-based base editing^8^. In the SMALT system, the AID/iSceI fused protein precisely targets a DNA barcode sequence (3kb in length), leading to cytosine –(C) - uracil (–U) - thymine (T) transitions through DNA replications^8^ (**Supplementary Fig 4**)). The original study provided cell phylogenetic trees for 16 organs from 2 fruit fly embryos (**Fig. 3a, b**). For scPhyloX analysis, nine organs were selected for the largest cell number in each fly (n>100 cells): brain disc (Br), eye-antennal discs (Ey), fat body (Fb), leg discs vT1 (L1), leg discs vT2 (L2), leg discs vT3 (L3), midgut (Mg), Malpighian tubule (Mp) and wing discs (Wg). We first estimated the barcoding mutation rate of each organ (**Supplementary Figs. 5, 6, Supplementary Table 2**). The estimated results indicate that the gene editing rate by the SMALT system during fruit fly development is approximately 0.5-1.5 mutations per cell division, which is consistent with the estimated values of the original study^8^ (**Fig. 3c**). Importantly, the mutation rate of the same organ shows high consistency between two flies (Spearman’s *ρ* = 0.67, *p* = 0.025) (**Fig. 3d**).

**Fig. 3.**
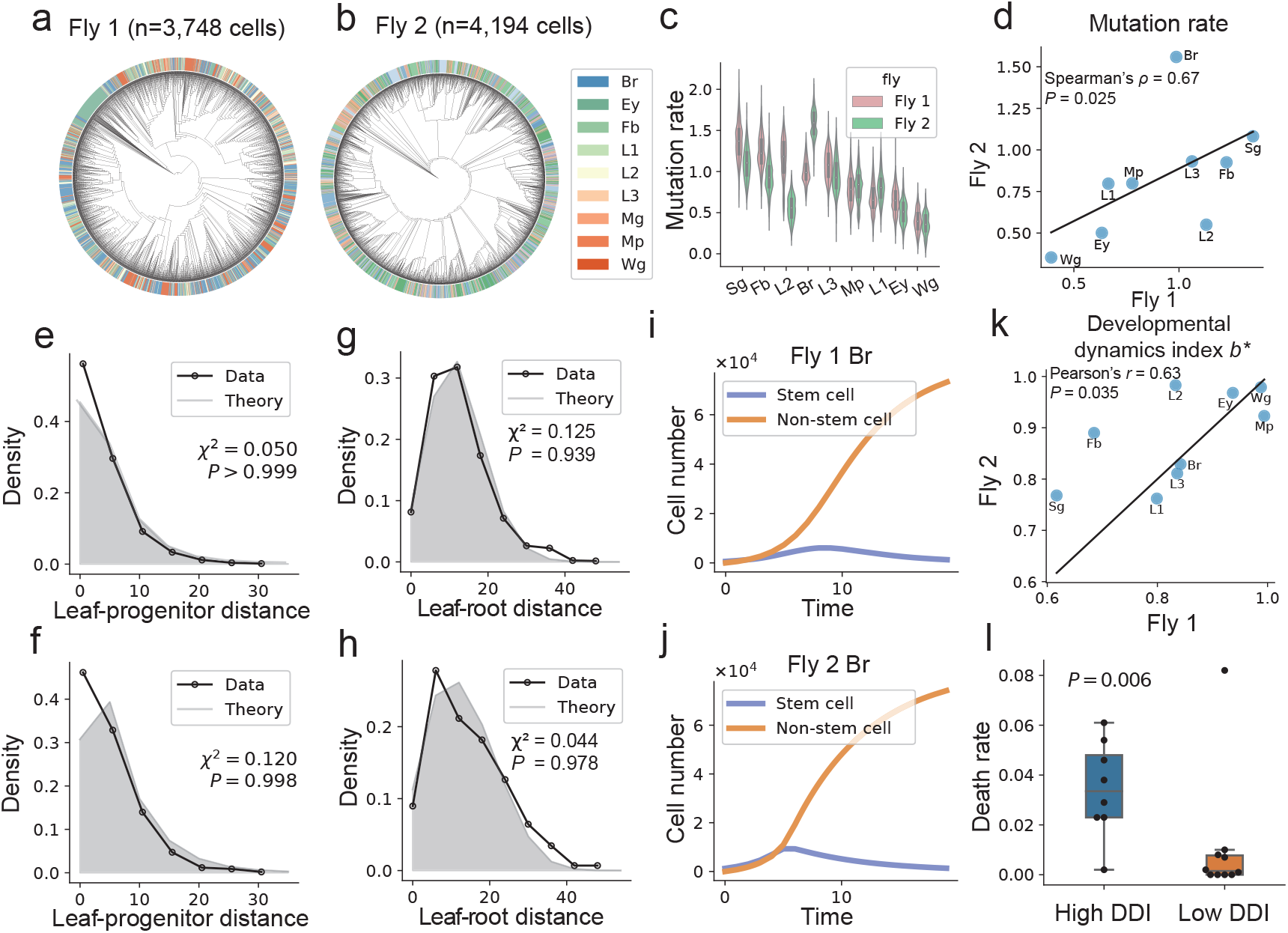
scPhyloX identifies overshoot development of fruit fly embryo development. (**a-b**), Phylogenetic trees of single cells sampled from 9 organs of two fly embryos where colors represent organs. Data are from Liu et al ^8^ that used the SMALT lineage tracing system. (**c**), Estimation of the base editing rate (i.e. mutation rate) per cell division of the SMALT system. Comparison of mutation rates across 9 different organs of fly 1 (red) and fly 2 (green). (**d**), Correlation of mutation rates between fly 1 and fly 2 across different organs. Spearman correlation and *P* value are shown. (**e-f**), Model fitting of the distributions of leaf-progenitor distances for brain tissue (Br) from fly 1 (**e**) and 2 (**f**), respectively. (**g-h**), Model fitting of the distributions of leaf-root distances for brain tissue (Br) from fly 1 (**g**) and 2 (**h**), respectively. (**i-j**), The inferred population growth of stem cells and non-stem cells during development of brain in fly 1 and 2, respectively. (**k**),Correlation of developmental dynamics index (DDI) between fly1 and fly2 across different organs. Spearman correlation and *P* value are shown. (**l**), Boxplot showing estimated death rate *d* for organs between high DDI (*b*^*^ ≥ 0.85) and low DDI (*b*^*^ *<* 0.85). Mann-Whitney U test *P* value is shown here.

We then used the dynamics model given by **Eq. (3)** to estimate the growth curve of stem and non-stem cells (**Supplementary Fig. 7-12**). Taking the brain disc as an example, we found that our model-inferred distance distributions (LP and LR) matched the actual data well, where consistent developmental growth dynamics of the two fruit flies was recovered (**Fig. 3e-j**). In fact, the parameter inferences for growth of the 9 organs were highly consistent between two fruit fly individuals (DDI, Pearson’s *r* = 0.63, *p* = 0.035) (**Fig. 3k**). Interestingly, the inferred DDI were generally small (*b*^*^ < 1), indicating overall overshooting growth of the stem cells during fly organ development (**Supplementary Figs. 9 and 10, Supplementary Table 2, Methods**). We also noticed that brain disc (Br) showed an exceptional difference in gene editing rate between the two flies (**Fig. 3d**). However, the DDI for brain disc were highly consistent, suggesting that scPhyloX is robust to mutation rate variations.

We further investigated the impact of the overshoot model on fruit fly development. Although all organs in two flies are mathematically defined as overshooting growth, we also observed that some organs with DDI close to 1 did not have a significant decline in stem cells (**Supplementary Figs. 11 and 12**). Therefore, we classified the organs into two categories: high DDI (*b*^*^ > 0.85, including Wg, Ey, Mp in both flies; weak overshooting effect), and low DDI (*b*^*^ < 0.85, strong overshooting effect). A comparative analysis of death rates revealed organs with a low DDI exhibited a lower death rate of non-stem cells compared to those with high DDI (*P* = 0.006) (**Fig. 3l**). This indicates that stronger overshoot of development can achieve the same cellular composition with fewer divisions than continuous growth, while also diminishing heterogeneity in cell divisions, significantly enhancing developmental efficiency and reducing development time.

### The clonal expansion of multipotent progenitors of human hematopoiesis during aging

We next applied our model to the phylogenies of hematopoietic stem cells or multipotent progenitors (HSC/MPPs) from whole-genome sequencing (WGS) of single-cell-derived colonies across 10 human subjects spanning from 0 to 81 years of age^6^. Here, the two cell populations are HSCs and MPPs, corresponding to stem cells and non-stem cells in the model, respectively. Because more differentiated blood cell types produced after MPPs differentiation were incapable of *in vitro* cultures and thus have not been sampled in the phylogenetic tree, so the event of “cell death” (i.e., the death rate *d*) estimated by the model here corresponds to the differentiation rate of MPPs. We applied scPhyloX to 8 non-embryonic samples aged 29-81 years each including 315-451 single HSCs or MPPs. By fixing the final HSC population size as ∼100,000 cells according to the original study, we used scPhyloX to interrogate the developmental dynamics of HSCs and MPPs. Interestingly, all samples showed a continuous growth of HSCs (**Fig. 4a-f, Methods, Supplementary Figs. 13a, 14, 15, Supplementary Table 3**). Notably, the number of MPPs at plateau increased during aging (**Fig. 4e-g, Supplementary Fig. 14**), where there was there was a significant negative correlation between the proportion of HSCs and age (Pearson’s *r* = −0.75, *P* = 0.017, **Fig. 4h**). According to the phylogenetic tree, elevated clonal expansions occurred as increase of age (**Fig. 4a-b, Supplementary Fig. 14**). Indeed, the corrected Colless’ index *CCT*(*T*) of cell phylogeny showed a significant positive correlation with age (Pearson’s *r* = 0.79, *P* = 0.01, **Fig. 4i**), in line with more stringent clonal expansions in elderly individuals. There data together suggested that clonal hematopoiesis is likely driven by expansion of MPPs instead of HSCs.

**Fig. 4.**
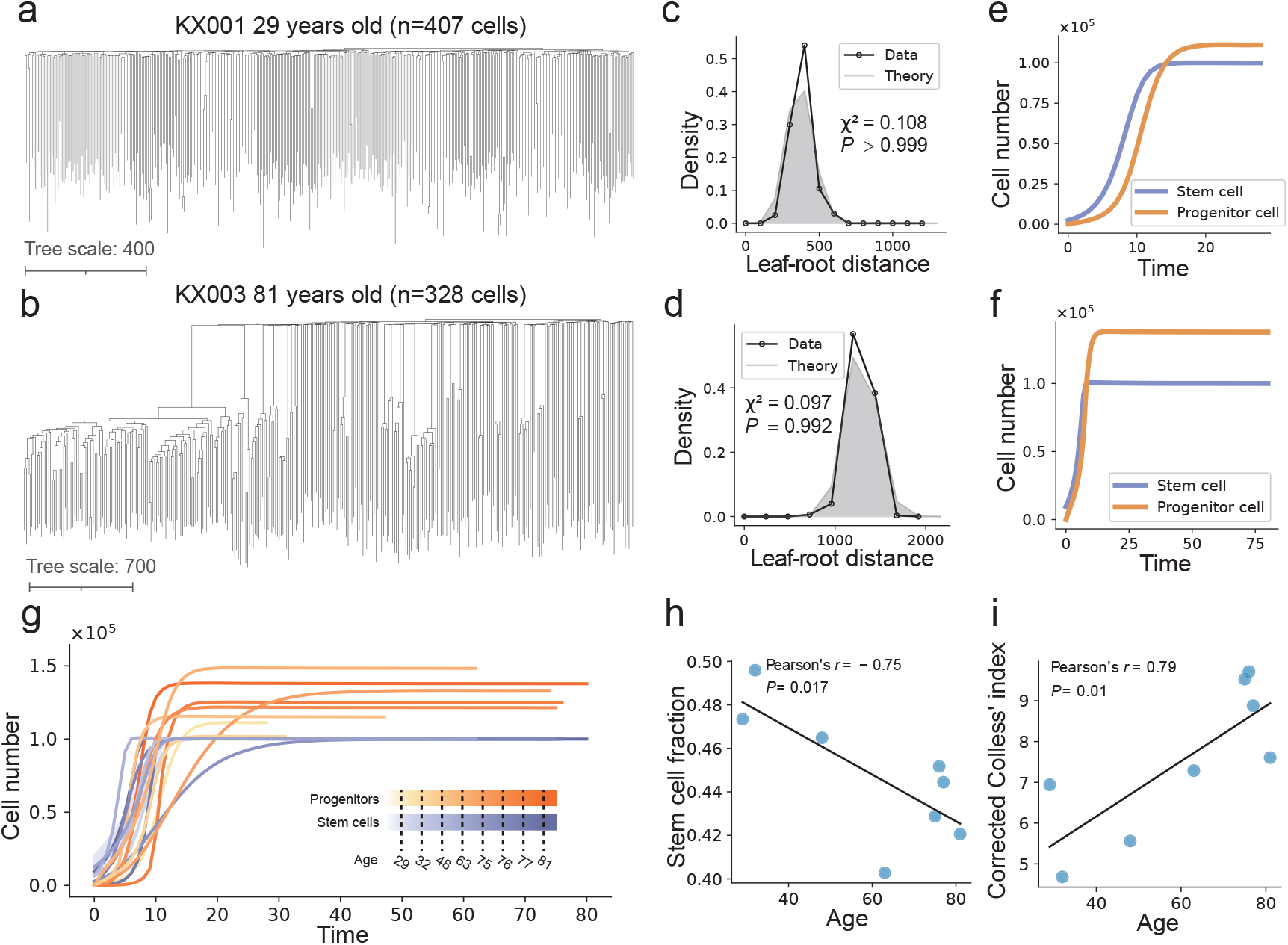
The cell population growth and clonal dynamics during human hematopoiesis. (**a-b**), Phylogenetic trees of 407 and 328 hematopoietic stem cells and multipotent progenitors (HSCs/MPPs) from a 29-year-old male (**a**) and 81-year-old male (**b**) donor, respectively. (**c-d**), Model fitting of the distribution of leaf-root distances in this two individuals, respectively. Pearson’s chi-square tests are shown. (**e-f**), The inferred cell population growth of stem cells (HSC) and progenitors (MPPs) in these two samples. (**g**), The inferred cell population growth of stem cells (HSC) and progenitors (MPPs) for all 8 samples across different ages. (**h**), Correlation between age and stem cell fraction. Pearson’s correlation and *P* value are shown. (**i**), Correlation between age and corrected Colless’ index of the tree. Pearson’s correlation and *P* value are shown.

### Strong subclonal selection in early colorectal tumorigenesis

We then extended scPhyloX to model tumor growth and subclonal selection, where the structured population consists of two types of cells: neutral founding cells and advantageous cells under positive subclonal selection. The growth rate of neutral cells is *ar*, while that of advantageous cells is (*a* + *s*)*r*, where *a* and *r* denotes the probability self-renewal of neutral cells and the rate of cell reaction, while *s* denotes a selective benefit. Moreover, a neutral cell acquires a beneficial driver mutation and becomes an advantageous cell with a probability *u* at each cell division. We also assumed a probability *d*_*NE*_ = 1 − *a* − *u* and *d*_*AD*_ = 1 − *a* − *s* that leads to cell death during cell division for neutral and advantageous cells, respectively (**Fig. 5a, b**). Let *NE* and *AD* denotes the numbers of neutral and advantageous cells, respectively. The cell number dynamic equations are given by

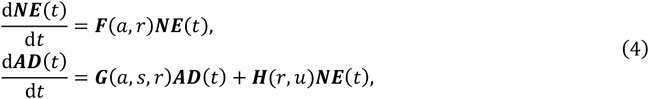

where ***N****E*(*t*) = (*NE*_0_(*t*), ⋯, *NE*_*n*_(*t*))^*T*^, ***AD***(*t*) = (*AD*_0_(*t*), ⋯, *AD*_*n*_(*t*))^*T*^, *NE*_*i*_ (*t*) and *AD*_*i*_ (*t*) are the numbers of neutral cells and advantageous cells in the i-th generation at time *t*, ***F, G*** and ***H*** are coefficient matrices of the equations.

**Fig. 5.**
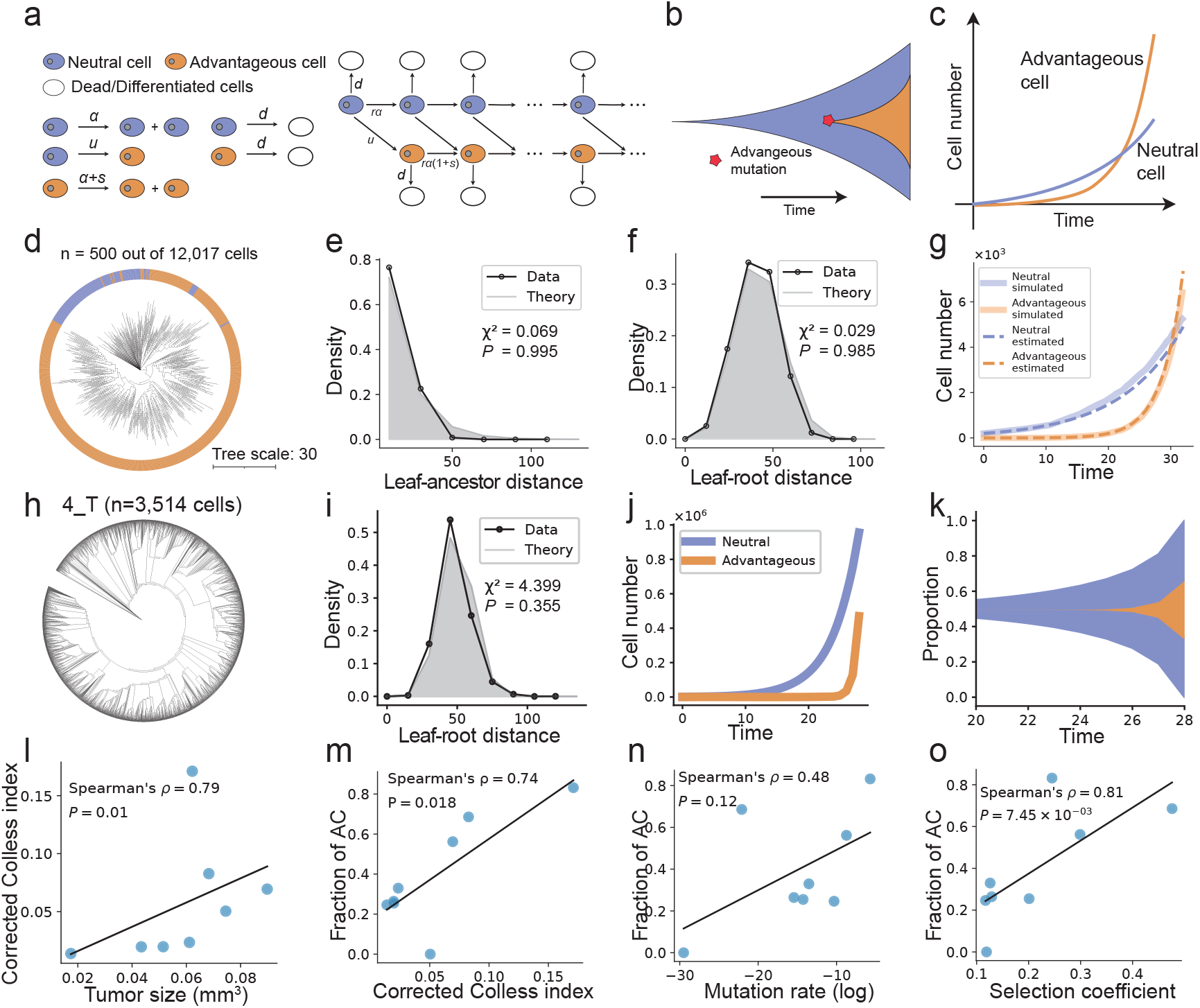
The evolutionary dynamics of neutral and advantageous cells in early colorectal tumorigenesis. (**a**), Cell-state evolution model of tumor growth. For a founder neutral cell (purple), it can divide into two neutral cells, become an advantageous cell (orange) through mutation or die at each cell cycle. For an advantageous cell (orange), it can divide into two advantageous cells or die, at each cell cycle. (**b**), Muller’s plot showing the emergence of advantageous mutations during tumor growth from a founder cell. (**c**), Growth of neutral (purple) and advantageous (orange) cells, where advantageous cells have a higher growth rate. (**d**), Phylogenetic tree of single cells sampled from simulated tumor. (**e**), Model fitting of the leaf-progenitor distance distribution. (**f**), Model fitting of the leaf-root distance distribution. Pearson’s chi-square tests are shown. (**g**), The inferred cell population growth (dashed lines) matches simulation results (solid lines) including both neutral (purple) and advantageous (orange) cells. (**h**), Phylogenetic tree of mouse colon tumor 4 T with 3,514 single cells. (**i**), Model fitting of leaf-root distance distribution for this tumor. Pearson’s chi-square tests are shown. (**j**), The inferred cell population growth of neutral (purple) and advantageous (orange) cells. (**k**), Muller plot showing the growth of neutral (purple) and advantageous (orange) cells. (**l**), Correlation between tumor size and corrected Colless index of the tree. Spearman’s correlation and *P* value are shown. (**m**), Correlation between corrected Colless index and fraction of advantageous cells. Spearman’s correlation and *P* value are shown. (**n**), Correlation between advantageous mutation rate (log-scale) and fraction of advantageous cells. Spearman’s correlation and *P* value are shown. (**o**), Correlation between selection coefficient s and fraction of advantageous cells. Spearman’s correlation and *P* value are shown.

Here, the parameters *u* and *s* are of interest, which together underlie the evolutionary dynamics of subclonal selection. The mutation rate *u* represents the probability of a beneficial mutation occurring. The selection coefficient *s* reflects the comparative growth rate of advantageous cells. By estimating these two parameters, we can measure the evolutionary dynamics of a tumor quantitatively. (**Fig. 5c, Methods**)

To examine the inference performance of tumor growth model, we simulated the process of tumor cells growing from neutral cells until the population size reached ∼10^4^. We set a certain probability of generating advantageous mutations during cell division (**Fig. 5d, Supplementary Fig. 16a**). Then, 500 cells were randomly sampled as sequenced cells, and their LR and LP distances were calculated for model inference. The results showed that scPhyloX can accurately infer the expansion of neutral and advantageous cells, as well as other model parameters including *λ, r, a, s* and *u* (**Fig. 5e-g, Supplementary Fig. 16b-e, Supplementary Table 1**).

We then applied scPhyloX to lineage tracing data of early colorectal lesions from mouse models of inflammatory-driven CRCs, recorded by the SMALT lineage tracing system^37^. We analyzed 8 mouse CRC samples (each including 126–3,803 cells in the phylogenetic tree) with uni-ancetral origin from the original study and inferred the growth dynamics of neutral and advantageous cells during tumor progression (**Fig. 5h-k, Supplementary Fig. 17-18, Supplementary Table 4**). We observed that the proportion of cells in advantageous clones, which have a selective advantage, varied among samples and often emerged relatively late in the tumorigenesis process. In 3 out of 8 samples (49_T1, 65_T1 and 66_T1), the advantageous cells in ensemble outnumbered neutral cells. The difference in proliferating fitness resulted in different degrees of imbalance of the cell phylogenetic trees. Interestingly, we discovered a significant positive correlation between corrected Colless index (CCI) of phylogenetic tree and tumor volume (Spearman’s *ρ* = 0.79, *P* = 0.010, **Fig. 5l**), as well as the proportion of advantageous cells (Spearman’s *ρ* = 0.74, *P* = 0.018) (**Fig. 5m**). It is posited that the advantageous clones within a tumor foster tumor growth and contribute to enhanced intratumoral driver-gene heterogeneity, in consistent with the branching evolution ^38, 39^.

Subsequently, we investigated the factors that facilitate the proliferation of advantageous cells. According to **Eq. (4)**, two parameters - the beneficial mutation rate *u* and the selection advantage *s* determine the proportion of the final advantageous cells. We found a weak correlation between the mutation rate *u* (log scale) and the proportion of advantageous cells (Spearman’s *ρ* = 0.48, *P* = 0.12, **Fig. 5n**). However, a significant correlation between selection coefficient *ss* and the proportion of advantageous cells was noted (Spearman’s *ρ* = 0.81, *P* = 0.007, **Fig. 5o**). This suggests that selection, rather than advantageous mutation rate, drives subclonal expansions in early colorectal tumorigenesis.

## Discussion

Phylodynamics is a powerful quantitative technique for inferring population dynamics from an observed phylogenetic tree^15, 19, 40^. Although previous studies have successfully applied phylodynamics models at the cellular level to infer population dynamics, there is still a lack of a general computational framework that models multiple cell types in somatic tissues based on single-cell lineage tracing data^21, 23, 24^. The novel framework we introduce in this study, scPhyloX, advances the field by providing a modeling framework of structured cell populations for quantitative phylodynamics inference of cell population dynamics. scPhyloX tackles complex scenarios involving multiple cell types, time-varying parameters, and somatic clonal evolution. Moreover, scPhyloX can reconstruct the natural history of tissue development and tumor evolution using single-time-point lineage tracing data, eliminating the need for time-series data or prior knowledge on cell phenotypes.

Analysis of lineage tracing datasets with scPhyloX across fly embryo development, human hematopoietic stem cells/progenitors and mouse colorectal tumors yields insights into termporal dynamics of cell populations. First, during the development of fruit fly embryos, the stem cells of each organ follow an overshooting growth pattern, which can significantly reduce the cell death rate during development and reduce the accumulation of deleterious mutations. In this regard, the overshooting might be an important mechanism of developmental robustness. Second, in human hematopoiesis, HSCs proliferate in a continuous growth pattern and the proportion of HSCs declines with aging. Finally, during the growth of mouse colorectal tumors, we found a scenario of low driver mutation rate but each driver confers a high selective fitness advantage, indicating selection rather mutations drive the subclonal expansions in early tumorigenesis.

Despite the rapid development of gene editing-based lineage tracing methods and their wide applications ^8, 36, 41, 42^, quantitative methods for analyzing cell lineage tracing datasets are still lacking, which is a bottleneck in this field. We believe scPhyloX provides a framework for tackling this problem and also stimulates the development of new methods that consider more sophisticated scenarios of tissue development and cancer evolution, such as the integration of single-cell RNA-seq datasets for delineating cell-state transition and evolution.

In summary, we provide a theoretical framework for quantitative phylodynamics inference using single-cell lineage tracing data. With the rapid emergence of high-quality single-cell phylogenetic data technologies, we expect scPhyloX to reveal new biological processes, including exploring developmental dynamics, tissue design principles and disease progression.

## Methods

### The expected distribution of leaf-progenitor (LP) distance

We assume that the number of mutations required by a cell in a single division follows a Poisson distribution

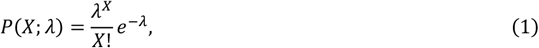

where *λ* is the mutation rate (measured in total mutation per cell division). Based on the properties of Poisson distribution, the number of novel mutations accumulated by a cell after *n* divisions also follows a Poisson distribution:

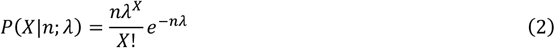

Regarding the leaf-progenitor (LP) distance, we must account for cell death or differentiation, which leads to lineage loss. We denote *δ* is the probability of death/differentiation per cell division. The number of cell divisions in last branches of phylogenetic tree *m* follows a Geometric distribution

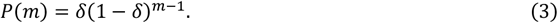

Then, we can calculate the length of the last branch in phylogenetic trees, which we refer to as the LP distance distribution

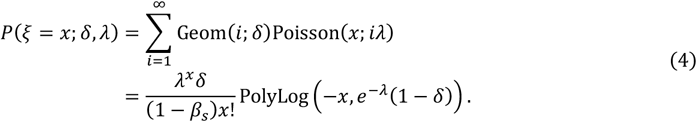

### Dynamic model of hierarchical tissue development

Assuming that we observe a tissue composed of *n* cells, which consists of two types of cells: stem cells (*SC*) and non-stem cells (*NC*). The stem cells perform cell division and tissue expansion functions, while the non-stem cells cannot continue to divide. We use a continuous-time Markov process to model the cell division process and denote the number of stem cells of generation *k* as *SC*_*k*_, and the number of non-stem cells of generation *kk* as *NC*_*k*_. Then, the state space of the Markov process is the vector of cell numbers of each generation of the two types of cells (*SC*_0_, *SC*_1_, ⋯, *SC*_*n*1_, *NC*_1_, *NC*_2_, ⋯ *NC*_*n*2_).

We assume that stem cell division has two modes, one is self-renewal, which produces stem cell with probability *β*(*t*), and cell differentiation, which produces non-stem cells with probability 1 − *β*(*t*). Differentiation can occur symmetrically or asymmetrically. Symmetric differentiation produces two non-stem cells with probability *p*, while asymmetric differentiation produces one stem cell and one non-stem cell with probability 1 − *p*. For non-stem cells, we assume that their death rate is *d*. Thus, we have

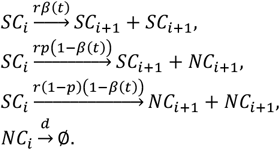

We can easily derive the expectation number of stem cells and non-stem cells using the following equations

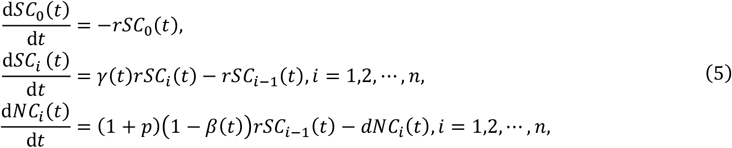

where *γ*(*t*) = 1 + *β*(*t*) − *p*(1 − *β*(*t*)).

Based on the fact that normal embryos do not grow indefinitely but grow to a fixed size and remain unchanged, we assume that the probability of stem cell division monotonically decreases and its derivate converges to 0. Therefore, we choose a sigmoid function

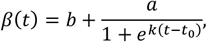

where *b* ∈ [0,1], *a* ∈ [0,1]. Then, *γ*(*t*) could be written as

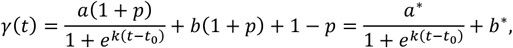

where *a*(1 + *p*) ≜ *a*^*^ ∈ [0,2], *b*(1 + *p*) + 1 − *p* ≜ *b*^*^ ∈ [0,2], *a*^*^ + *b*^*^ ≤ 2. Then, we can solve the equation as follows:

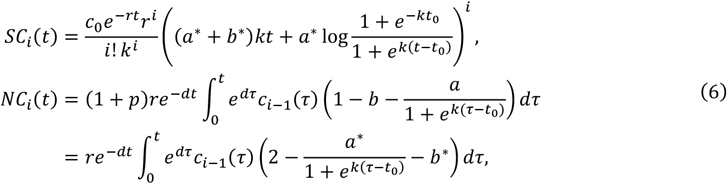

The parameter *b*^*^ plays a crucial role in the dynamics of stem cells. When *b*^*^ ≥ 1, the number of stem cells grows continuously, a phenomenon we refer to as continuous growth. When *b*^*^ < 1, the number of stem cells first increase and then decreases, a phenomenon we refer to as overshoot.

### Dynamic model of subclonal selection in tumor growth

Regarding the growth process of tumors, we focus on studying the evolutionary relationship between tumor neutral cells and advantageous cells. We assume a tumor is composed of neutral cells and advantageous cells. The neutral cells are normal tumor cells with growth rate *ar*, and the advantageous cells are tumor cells with positive selection, with growth rate (*a* + *s*)*r*. Moreover, there is a probability *u* that a neutral cell acquires a positive mutation and becomes an advantageous cell.

There is also a probability *d*_*NE*_ and *d*_*AD*_ that leads to cell death during cell division for both neutral and advantageous cells. Similarly, let *NE* and *AD* denote the numbers of neutral and advantageous cells, respectively. Then we have

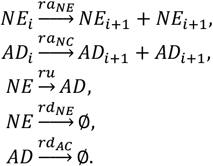

Without loss of generality, we assume *a*_*AD*_ = *a*_*NE*_ + *s*. Notice that *a*_*NE*_ + *u* + *d*_*NE*_ = 1, *a*_*AD*_ + *d*_*AD*_ = 1, the cell number dynamic equations are given by

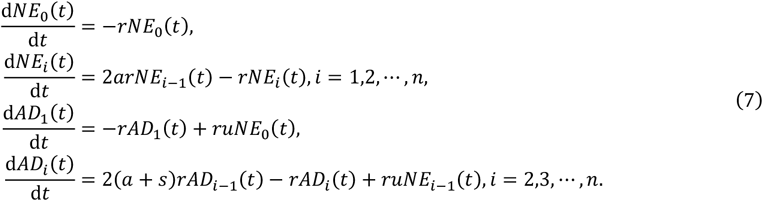

Then, we can solve the equation using numerical method, here we use the Runge-Kutta method of order 5(4) (RK45)^43^ method in Scipy^44^.

### Simulation of single-cell phylogenetic data

We employed continuous-time Markov processes to model the development of single cells into complete tissues. The Gillespie algorithm was utilized, requiring only the rates of cell division, differentiation, and death for simulation.

In the overshoot developmental model, we distinguished between stem and non-stem cells. The division rate of stem cells into two new stem cells was denoted as

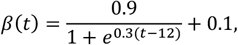

while the rate of symmetric differentiation into non-stem cells was *α*(*t*) = 0.6 (1 − *β*(*t*)), and the probability of asymmetric differentiation into one stem cell was *α*(*t*) = 0.4(1 − *β*(*t*)). The death rate of non-stem cells was set to 0.01.

In the continuous growth model, we distinguished between stem and non-stem cells. The division rate of stem cells into two new stem cells was denoted as

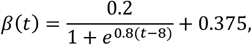

while the rate of symmetric differentiation into non-stem cells was *α*(*t*) = 0.6 (1 − *β*(*t*)), and the probability of asymmetric differentiation into one stem cell was *α*(*t*) = 0.4 (1 − *β*(*t*)). The death rate of non-stem cells was set to 0.2.

In the tumor growth model, the division rates of neutral and advantageous cells were defined as 0.6 and 0.8 respectively, with the mutation rate of neutral cells to advantageous set t 0.001. The death rates were set to 0.4 and 0.2 for neutral and advantageous cells, respectively. The simulation was halted when the time reached *T* = 35.

During cell division, DNA mutations were also simulated, with the number of new mutations per division following a Poisson distribution with an expected value of 2. Upon completion of the simulation, approximately 20,000 cells were generated, from which 500 were randomly selected for inference analysis in scPhyloX.

### Parameter estimation of mutation rate

We first use **Eq. (4)** to estimate the mutation rate. Noting that **Eq. (4)** is determined by two parameters, the mutation rate *λ* and the branching division probability *δ*, we use the DEMetropolis method to sample the posterior distributions of these two parameters. Since *δ* ∈ [0,1], we use the Beta distribution as the prior distribution for the parameter *δ*:

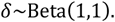

For the mutation rate *λ*, we first estimate the approximate mutation rate *λ*_0_ based on the number of mutations and the approximate number of cell divisions, and then use the truncated normal distribution as the prior distribution

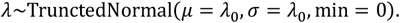

The proposal distribution of *δ* and *λ* are both normal distribution as default set in PyMC^45^.

### Calculate LR and LP distances from real data

For data in phylogenetic tree format (such as newick files), we can directly calculate the leaf-root (LR) distance and leaf-progenitor (LP) distance based on the tree structure. In practice, we first use biopython^46^ to read the phylogenetic tree file. For leaf-root distance, the *depths()* attribute is used to determine the distance from each leaf node to the root node. For LP distance, the *get_terminals()* attribute is used to traverse each leaf node and then calculate its distance from the nearest inner node.

For genome sequencing data, in addition to using tree reconstruction algorithms (such as maximum parsimony, maximum likelihood, etc.) to reconstruct its phylogenetic tree, we can also directly calculate LR distance and LP distance based on its mutation information. For LR distance, we count the number of mutation differences between the sequence of each cell and the reference sequence. For LP distance, we first calculate the Levenshtein distance matrix between all cells (here we use the XOR operation for equal-length sequences), and then find the minimum value of each row of the matrix except the diagonal.

### Estimation of the number of cell generations

We use maximum a posteriori estimation (MAP) to estimate the number of cell generations based on the LR distance (*η*) and mutation rate (*λ*). First, we know that for a given number of generations (*X* = *x*) and mutation rate (*λ*_1_), the distribution of LR distance is given by

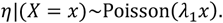

We approximate the distribution of the LR distance with a normal distribution, and approximate the prior distribution of the number of generations based on the mutation rate *λ*_1_ inferred in the previous step,

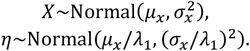

Then we have the estimation of the generation number *n* as follows,

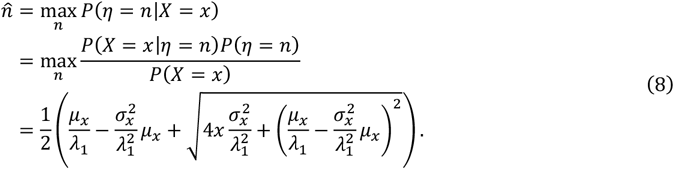

### Parameter estimation for phylodynamics inference

Next, we introduce the parameter estimation methods of dynamic equations (5), (7). Having derived the analytical expressions of these equations and estimated the number of cells in each generation, we use the Markov chain Monte Carlo (MCMC) method to estimate the posterior distribution of each parameter. We define the likelihood function as follows:

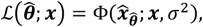

Where Φ is the probability density function of Normal distribution, with expectation ***x*** and variation *σ*^2^, ***x*** = (*x*_0_, *x*_1_, ⋯, *x*_*n*_)^T^ = (*NC*_0_ + *SC*_0_, *NC*_1_ + *SC*_1_, ⋯, *NC*_*n*_ + *SC*_*n*_)^T^ is cell number in each generation (or *x* = (*NE*_0_ + *AD*_0_, *NE*_1_ + *AD*_1_, ⋯, *NE*_*n*_ + *AD*_*n*_)^*T*^ for tumor growth model), 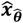 is the analytical solution of equation (5) (or equation (7) for tumor growth model) under 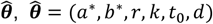 for tissue model, 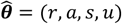 for tumor growth model.

Because our model is relatively complex and has many parameters, obtaining the global optimal estimates of the parameters directly using MCMC or other optimization methods is difficult. Therefore, we first search for the approximate global optimal points using the Differential Evolutionary algorithm (DE). Based on the practical significance of the parameters, we set the search range of each parameter as follows. For tissue development model,

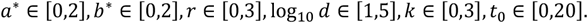

For tumor growth model,

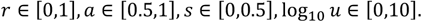

After obtaining the optimal differential evolutionary estimate 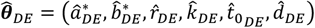, we use the DE Metropolis algorithm to sample the posterior distributions of the parameters. We set the prior distributions of each parameter as follows. For tissue development model,

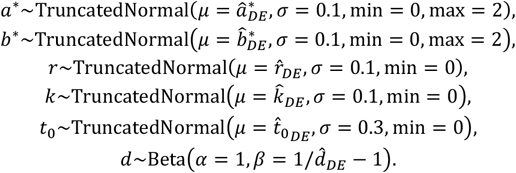

For tumor model,

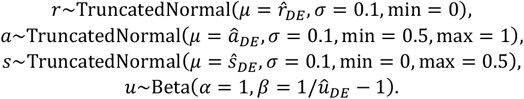

The proposal distributions of all parameters are set to normal distribution as default setting in PyMC^45^.

### Analysis of lineage tracing datasets

#### Datasets and pre-processing

We applied scPhyloX to four real lineage tracing datasets that are publicly available through online sources (‘Data availability’). These datasets include the in vitro culture of the human kidney cell line HEK293T^36^, SMALT recorded fruit fly embryos^8^, human HSC/MPPs^6^ and SMALT recorded mouse colorectal tumors^37^. The lineage trees of all four datasets were obtained from the original studies. For the fruit fly embryonic development dataset, we selected organs that are common to 2 flies and have more than 100 cells for analysis to ensure the accuracy of the inference results. In the human HSC/MPPs dataset, two embryo samples were excluded because their hematopoietic systems were not fully developed. In the mouse colorectal tumors dataset, 8 uni-ancestral tumors were analyzed for there are two types of cells fitted model assumption. All lineage trees were read, and branch lengths were calculated using biopython ^46^ and ete3 ^47^ and visualized using iTOL^48^.

#### Applying scPhyloX

For the human kidney cell line HEK293T dataset, we used a mutation rate *λλ* = 3 pre cell division from original study. All parameters in the tissue development model were estimated. For the fly embryo development dataset, **Eq. (1)** was used to estimate the mutation rate of each organ. Then, all parameters in the tissue development model were estimated. In the human HSC/MPPs development dataset, we first implemented the tissue development model with full parameters. We noticed that the estimated *b*^*^ in the 8 samples were around 1 (ranging from 0.95 to 1.01), and all showed a pattern of continuous growth of HSC (**Supplementary Fig. 13a**). To study the dynamic behavior of HSCs and MPPs more accurately, we fixed *b*^*^ = 1, i.e., the number of HSCs remained constant after they reached a certain population size (according to the original study, we had HSCs at a size of 100,000 cells). In this way, there was no significant difference between the log-likelihood of the 8-sample estimate and the log-likelihood of the flexible model with non-fixed *b*^*^ (**Supplementary Fig. 13b**). For the mouse colorectal tumors dataset, we used a mutation rate *λ* = 0.4 bp pre cell division from the original study. All parameters in tumor growth model were estimated.

### Quantifying the balance of phylogenetic tree

The Colless index, introduced by Colless D. H. ^49^, has been widely used in the analysis of phylogenetic tree. It quantifies the balance of a rooted bifurcating tree by considering the differences in subtree sizes induced by the children of inner vertices. For a rooted bifurcated tree *C*, the Colless index is given by

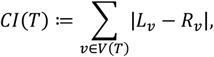

where *V*(*T*) is the set of all internal nodes in *T, L*_*v*_ and *R*_*v*_ are the number of left and right child nodes of node *v*, respectively.

Now, the corrected Colless index^50^ refines this concept. It normalizes the Colless index by adjusting for the number of leaves present in the tree. Essentially, it scales the pending subtree size differences induced by inner vertices, taking into account the overall leaf count. The corrected Colless index is defined as follows:

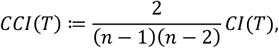

where *n* is number of leaf nodes of tree *C*.

## Data availability

All data analyzed in this article are publicly available through online sources. All lineage trees results and python implementation are available at https://github.com/kunwang34/scPhyloX. The raw data for HEK293T^36^ dataset can be accessed with PRJNA757179. The SMALT lineage tracing dataset for fruit fly embryos^8^ can be accessed with PRJNA716791. The dataset for human HSC/MPPs^6^ can be accessed with EGAD00001007851 and from Mendeley (https://data.mendeley.com/datasets/np54zjkvxr/1). The SMALT lineage tracing dataset for mouse CRC samples^37^ can be accessed from Zenodo (https://zenodo.org/records/10211904).

## Code availability

ScPhyloX is freely available as a Python package at https://github.com/kunwang34/scPhyloX. Detailed workflows to reproduce figures and results in this paper are available at https://scphylox.readthedocs.io/.

## Acknowledgements

We thank members of Zhou and Hu laboratories for constructive discussions. This work was supported by National Natural Science Foundation of China (11971405 to D.Z., 32270693 & 82241236 to Z.H.), Guangdong Basic and Applied Basic Research Foundation (2021B1515020042 to Z.H.) and Fundamental Research Funds for the Central Universities (20720230023 to D.Z.).

## Author contributions

K.W., Z.H. and D.Z. designed the study. K.W. developed the mathematical framework and implemented the software. K.W. analyzed the data. Z.L., Z.Y. and X.H. provided constructive suggestions on the methods. K.W., Z.H., D.Z., Z.L, Z.Y. and X.H. interpreted results. K.W., Z.H. and D.Z wrote the manuscript, with contributions from all co-authors. Z.H. and D.Z. supervised the study.

## Supplementary Information

Supplementary Figs. 1-18

Supplementary Tables. 1-4

**Supplementary Fig. 1.**
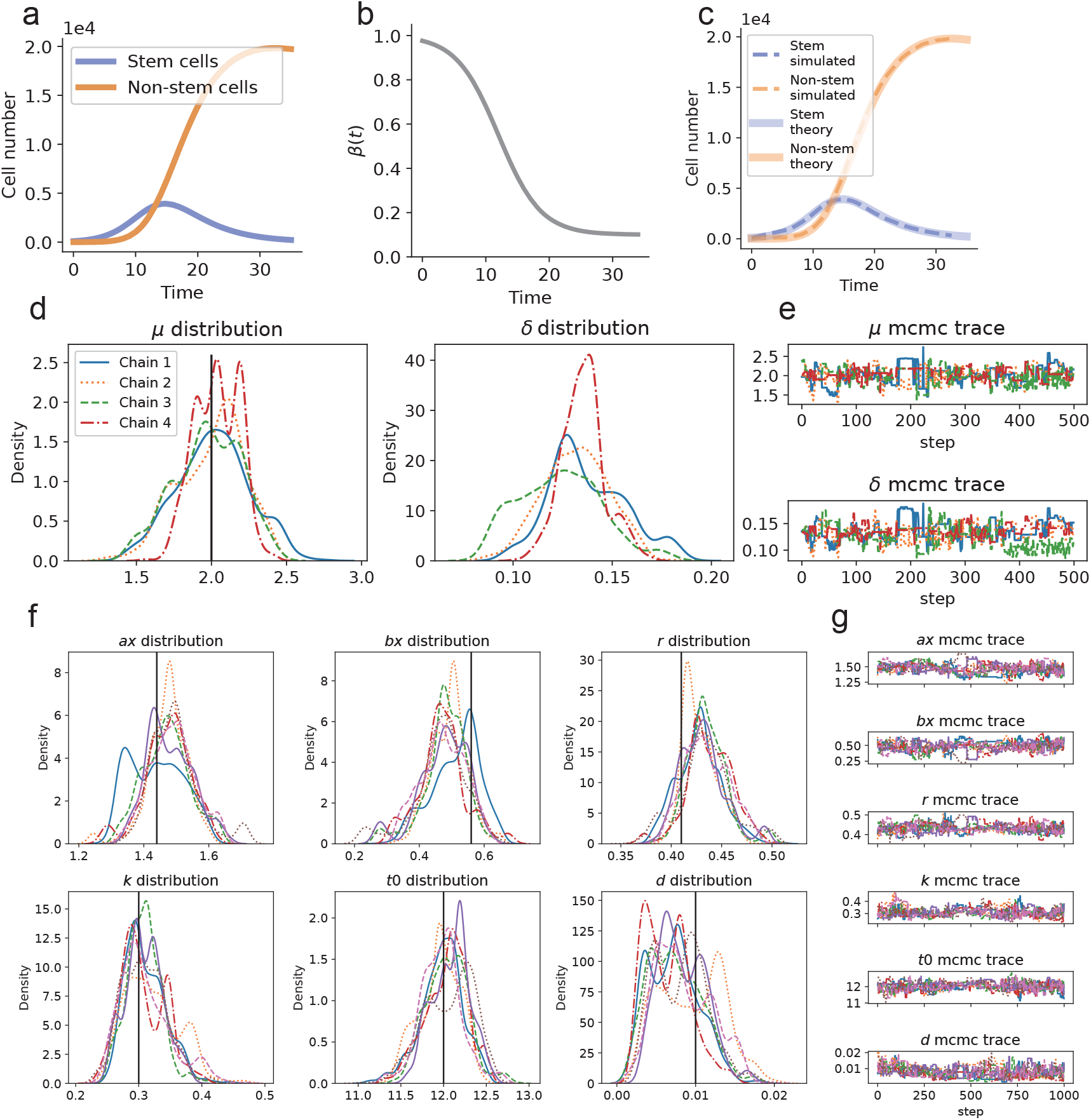
Overshoot model simulation results and parameter inference details. (**a**) Theoretical population growth under simulated overshoot model parameters. (**b**) The probability of stem cell replicated *β*(*t*) changes with time. (**c**) Theoretical population growth (solied lines) matches simulation (dashed lines). (**d**) Posterior distribution of mutation rate estimation. (**e**) MCMC sampling trace of mutation rate estimation. (**f**) Posterior distribution of phylodynamics parameters. (**g**) MCMC trace of phylodynamics parameters.

**Supplementary Fig. 2.**
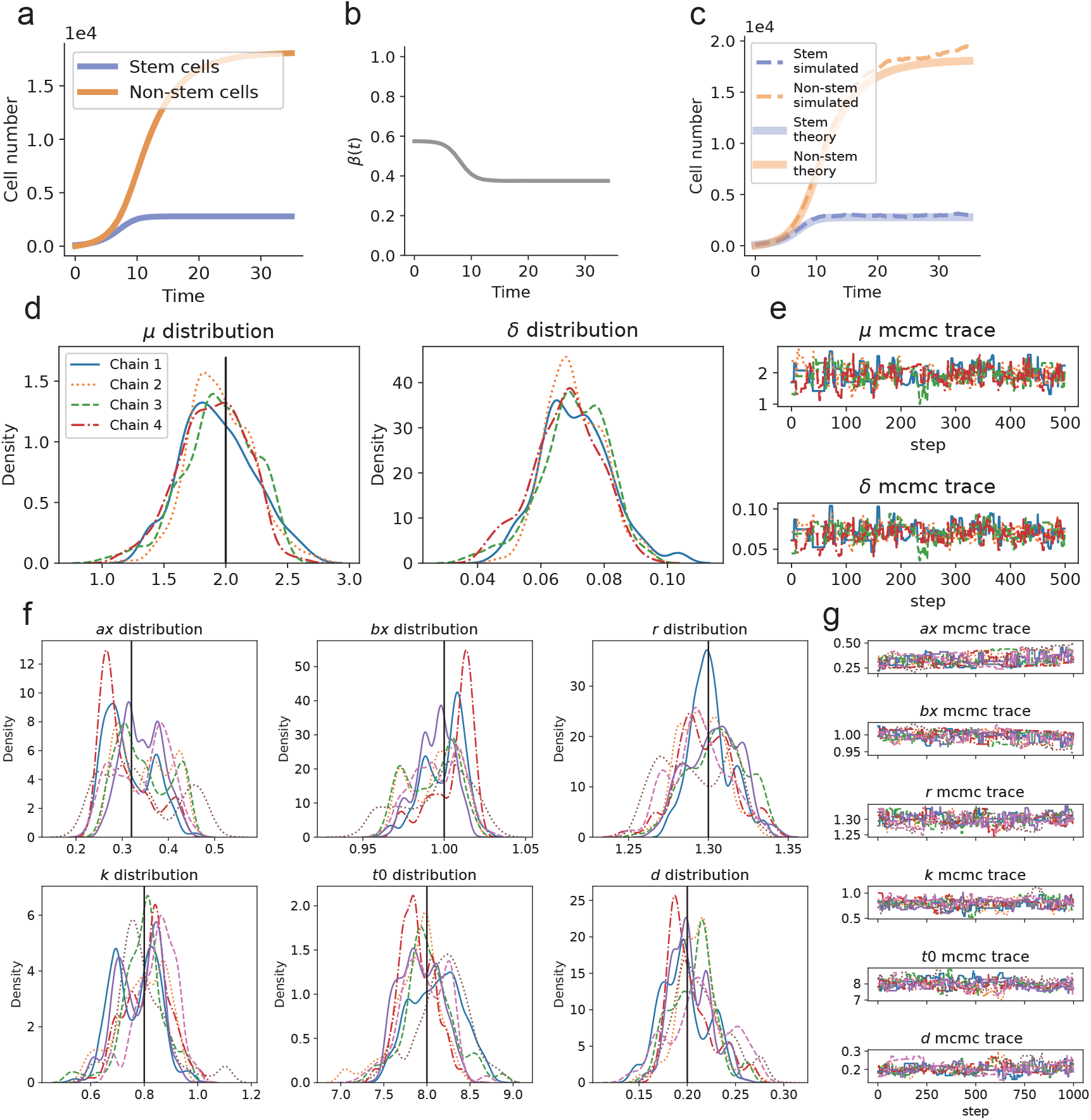
Countinuous growth model simulation results and parameter inference details. (**a**) Theoretical population growth under simulated continuous growth model parameters. (**b**) The probability of stem cell replicated *β*(*t*) changes with time. (**c**) Theoretical population growth (solied lines) matches simulation (dashed lines). (**d**) Posterior distribution of mutation rate estimation. (**e**) MCMC sampling trace of mutation rate estimation. (**f**) Posterior distribution of phylodynamics parameters. (**g**) MCMC trace of phylodynamics parameters.

**Supplementary Fig. 3.**
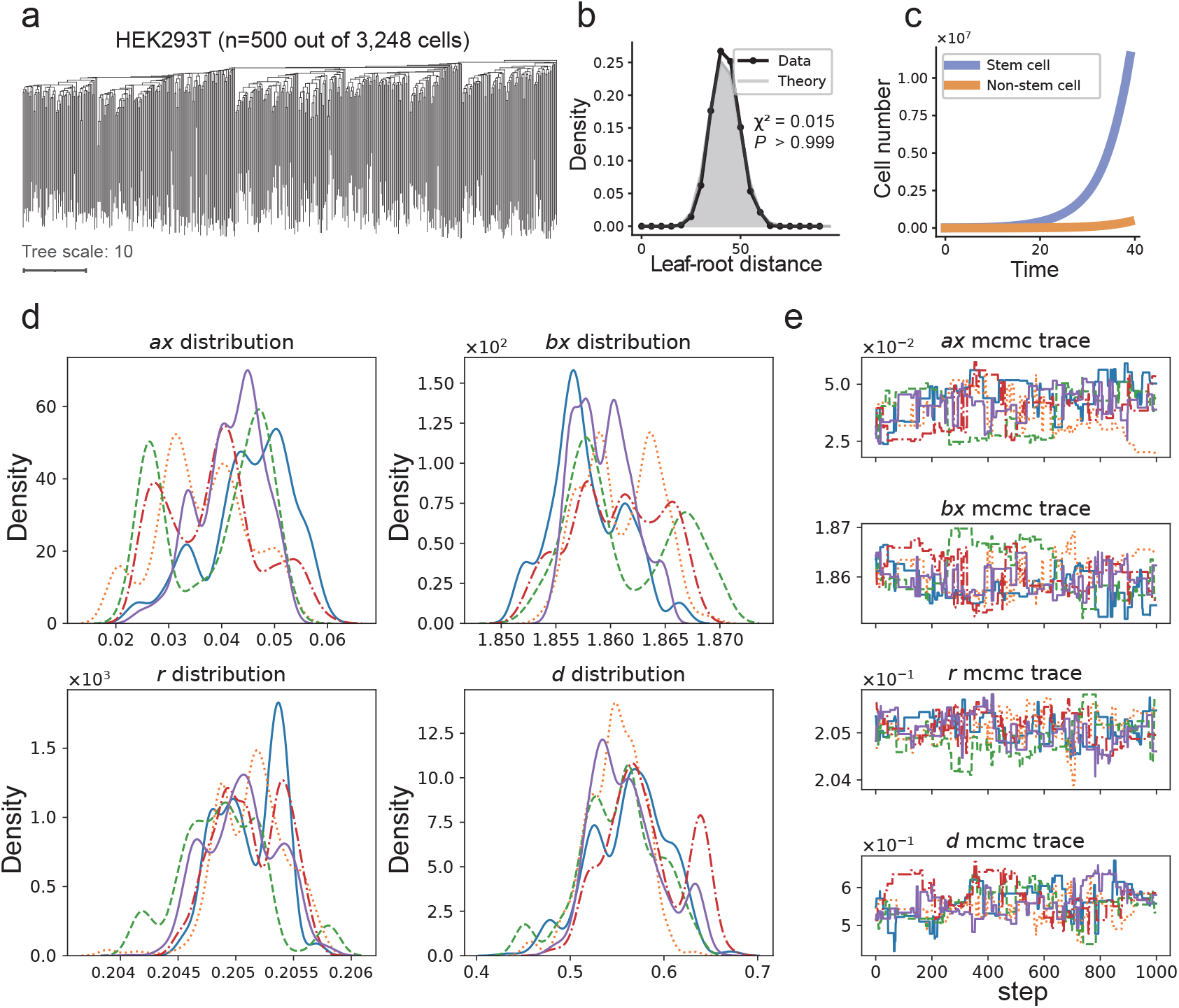
scPhyloX infers exponential growth of HEK293T cell line and parameter inference details. (**a**) Phylogenetic tree of 500 out of 3,248 HEK293T cells sampled from in-vitro culture of a clonal population. (**b**) Model fitting of the distribution of leaf-root distance. (**c**) The inferred population growth of stem cells and non-stem cells during development of HEK293T cells (**d**) Posterior distribution of phylodynamics parameters. (**e**) MCMC trace of phylodynamics parameters.

**Supplementary Fig. 4.**
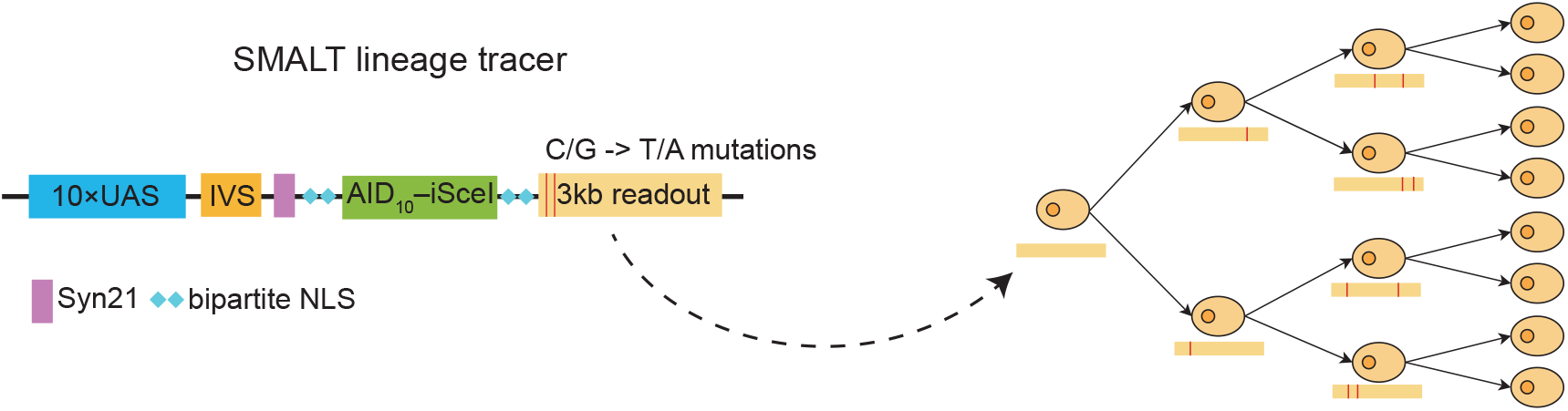
Schematic of SMALT lineage tracer system. SMALT records the history of cell division through AID-included single-nucleotide variation in developing DNA barcode.

**Supplementary Fig. 5.**
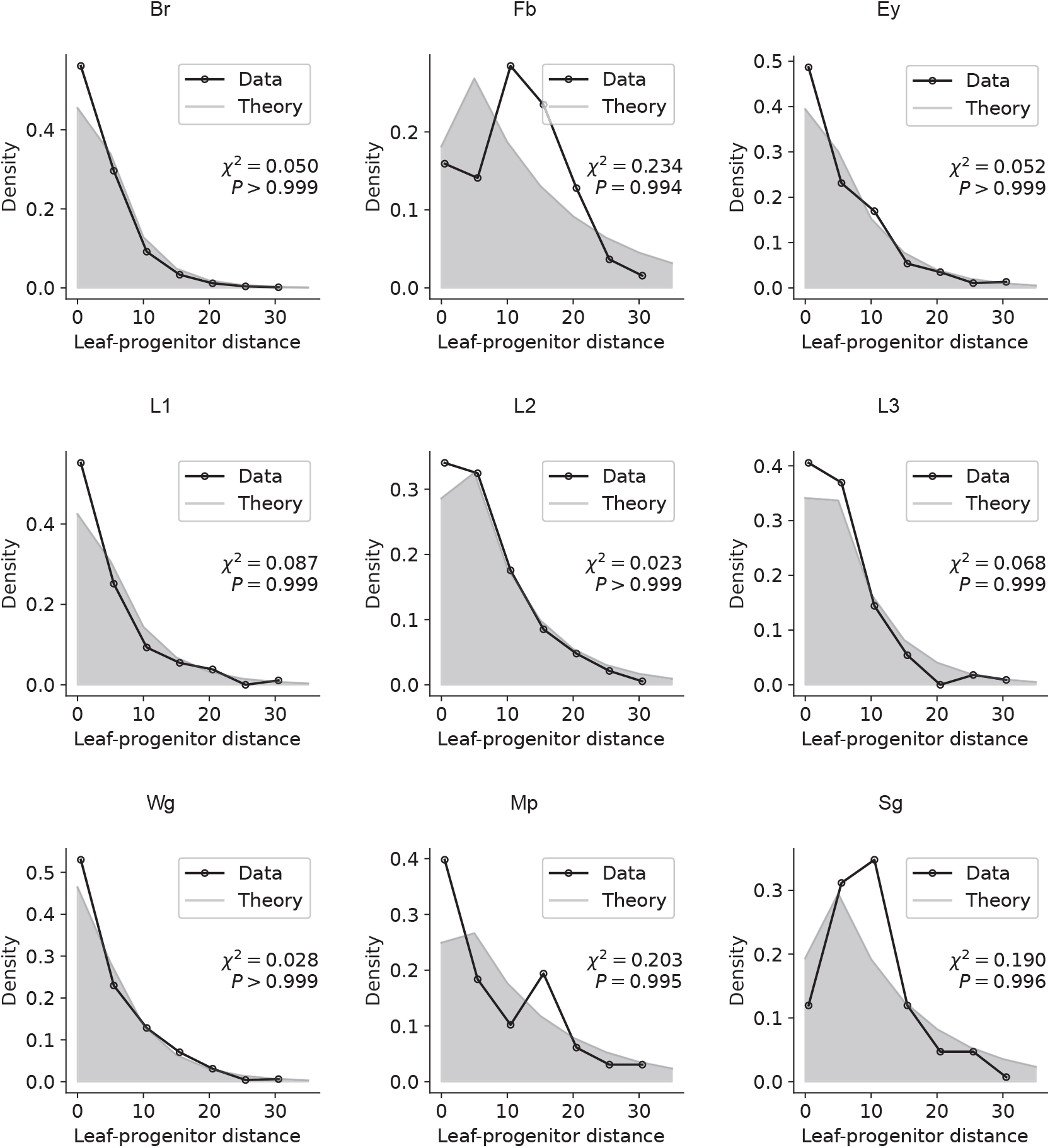
Model fitting of the distributions of leaf-progenitor distances for all tissues from fly 1. Leaf-progentior distributions for brain disc (Br), fat body (Fb), eye-antennal discs (Ey), leg discs vT1 (L1), leg discs vT2 (L2), leg discs vT3 (L3), midgut (Mg), Malpighian tubule (Mp) and wing discs (Wg) development. Pearson’s chi-square tests are shown here.

**Supplementary Fig. 6.**
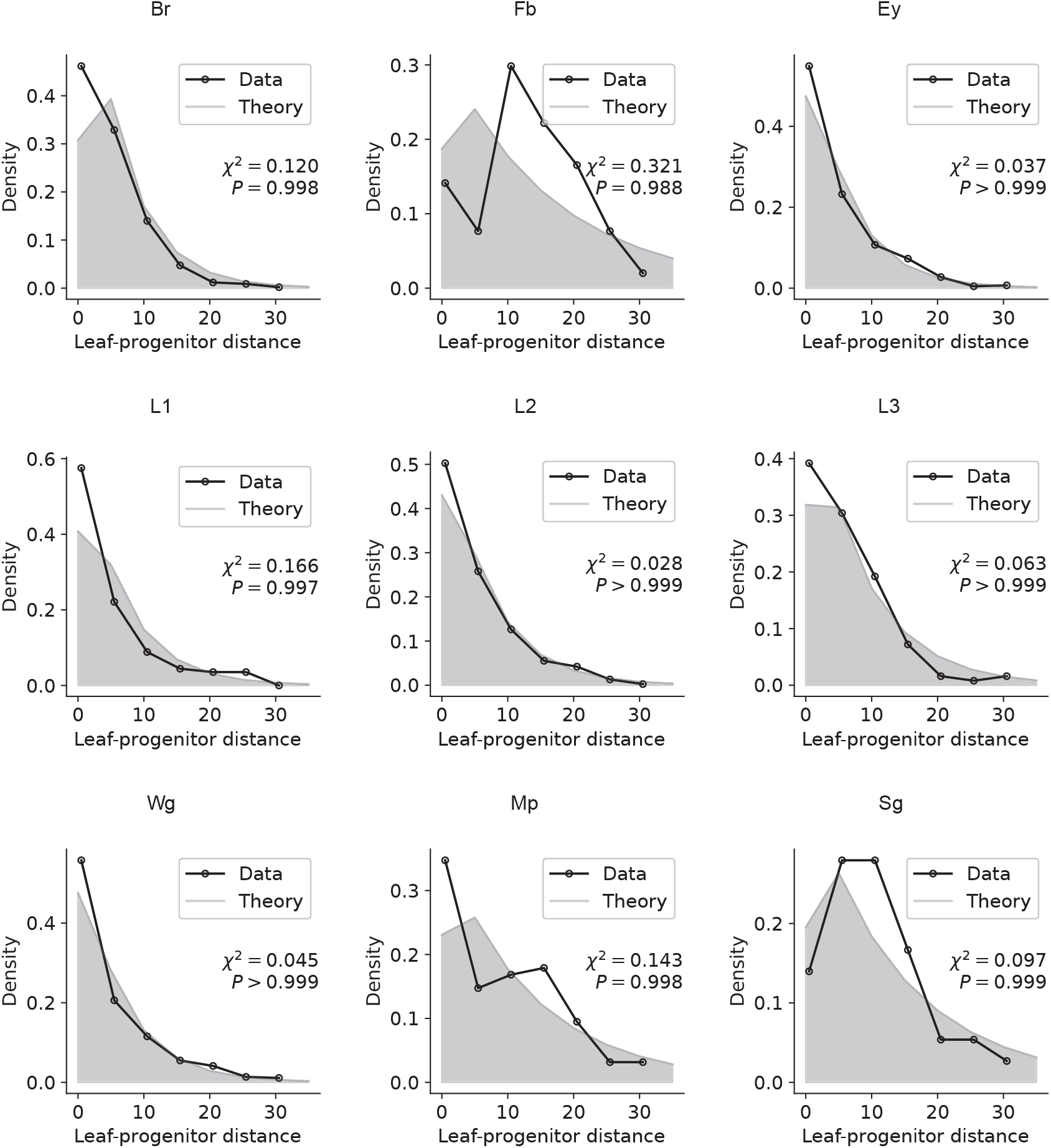
Model fitting of the distributions of leaf-progenitor distances for all tissues from fly 2. Pearson’s chi-square tests are shown here.

**Supplementary Fig. 7.**
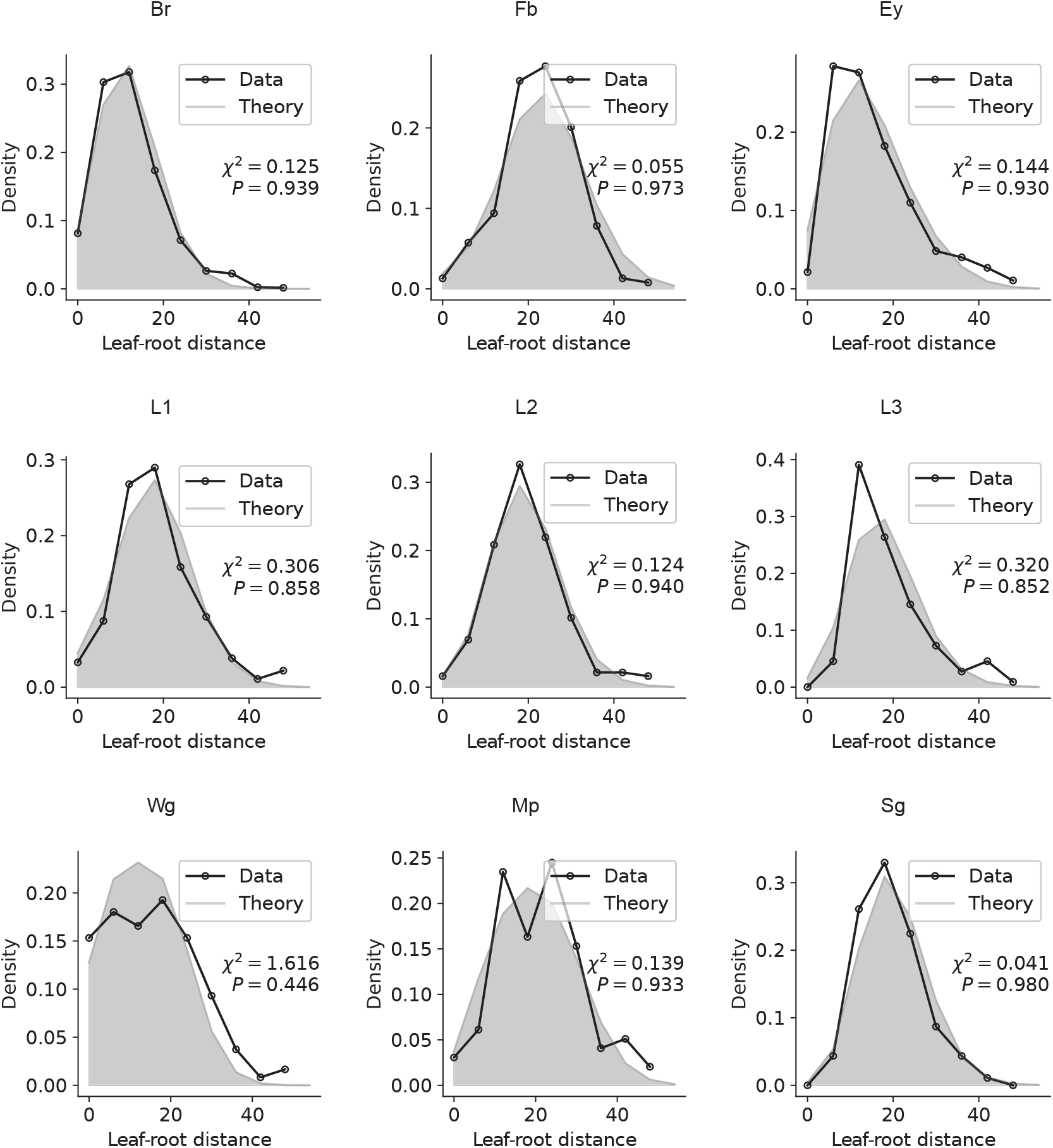
Model fitting of the distributions of leaf-root distances for all tissues from fly 1. Pearson’s chi-square tests are shown here.

**Supplementary Fig. 8.**
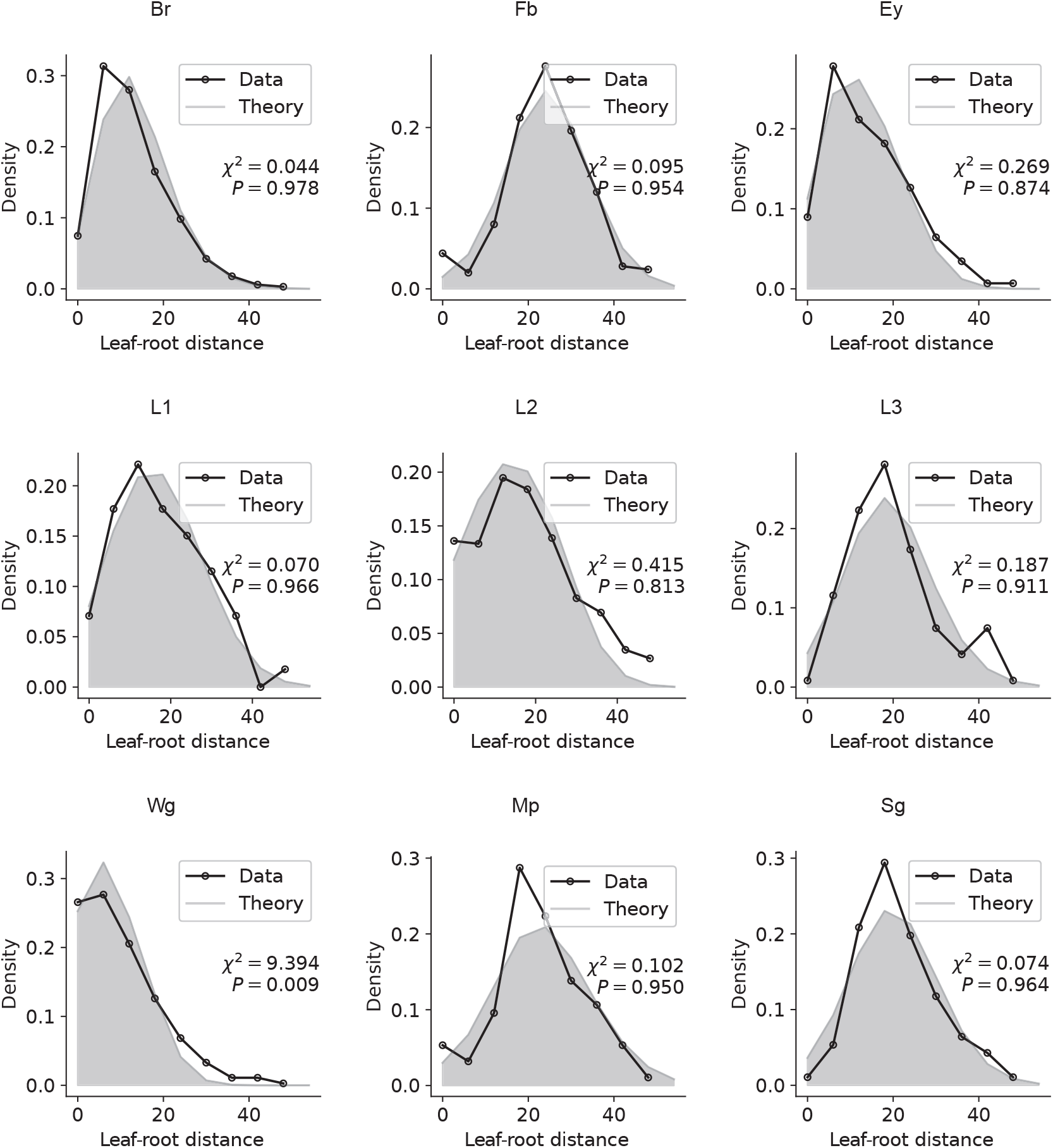
Model fitting of the distributions of leaf-root distances for all tissues from fly 2. Pearson’s chi-square tests are shown here.

**Supplementary Fig. 9.**
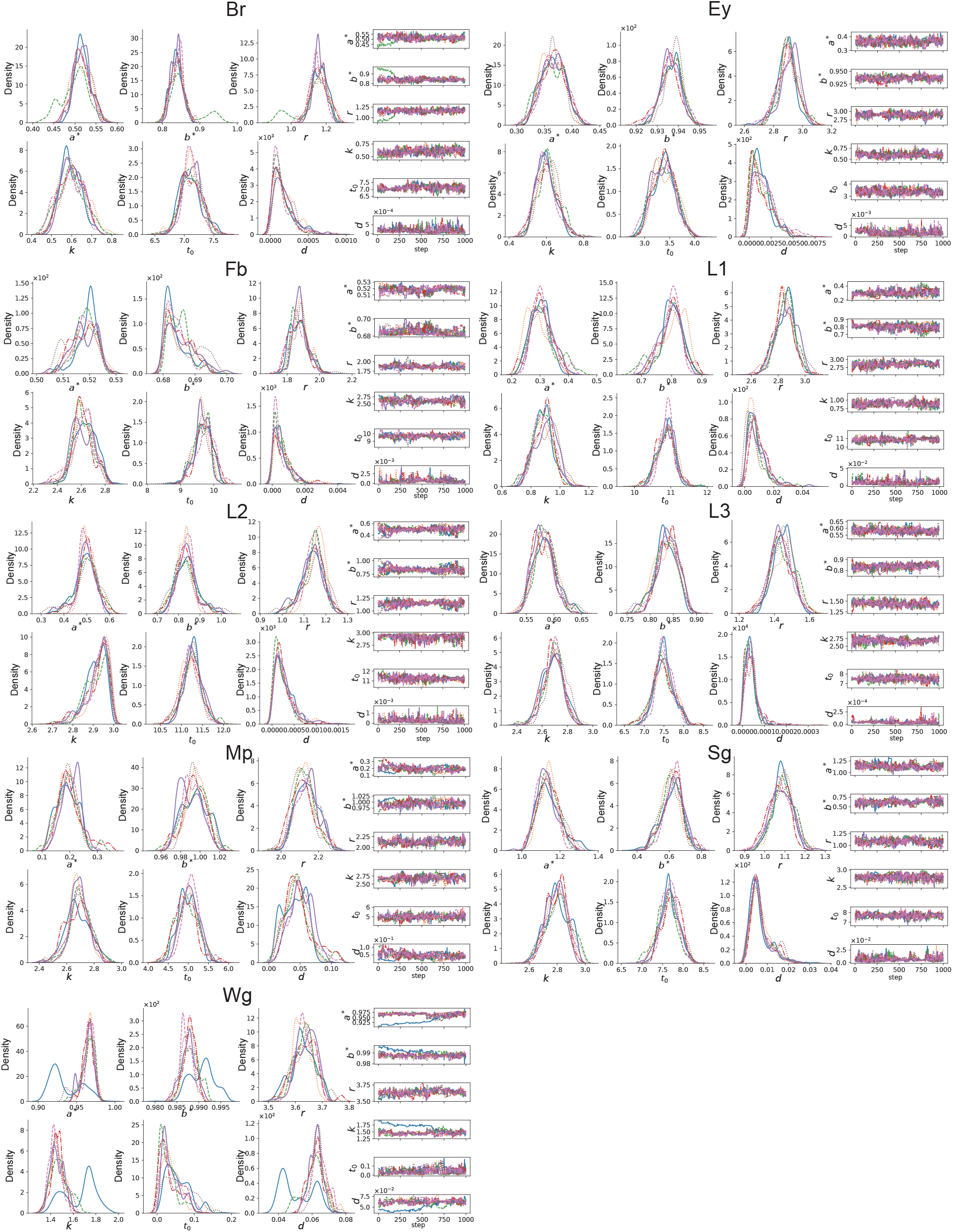
MCMC inference details of Fly1. MCMC inferred fly1 phylodynamics parameter distribution (left) and sampling trace (right) of each organ.

**Supplementary Fig. 10.**
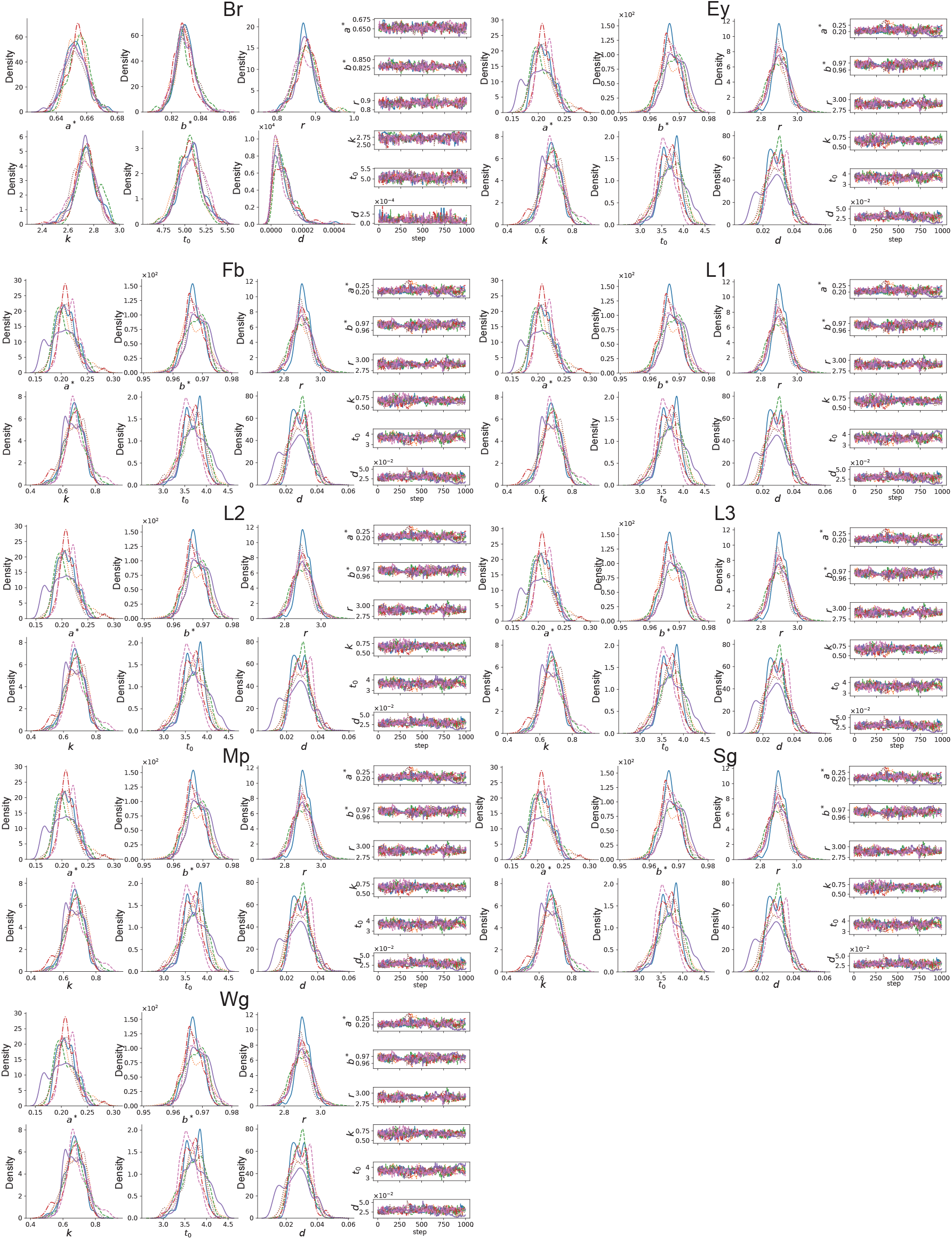
MCMC inference details of Fly2. MCMC inferred fly2 phylodynamics parameter distribution (left) and sampling trace (right) of each organ.

**Supplementary Fig. 11.**
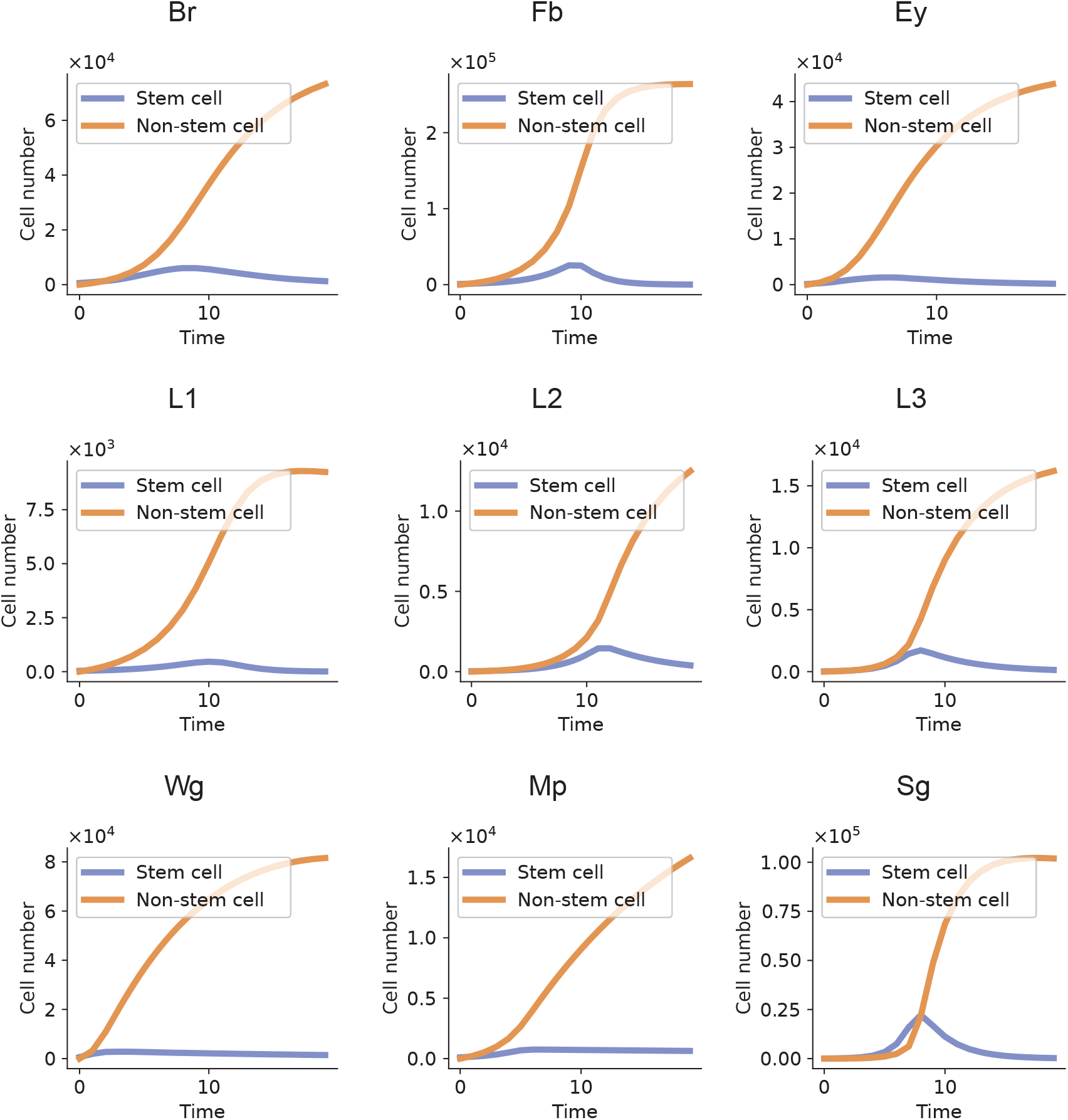
scPhyloX identifies overshoot development in Fly 1. The inferred population growth of stem cells and non-stem cells during development of fly 1. Stem cells and non-stem cells are marked in purple and orange, respectively.

**Supplementary Fig. 12.**
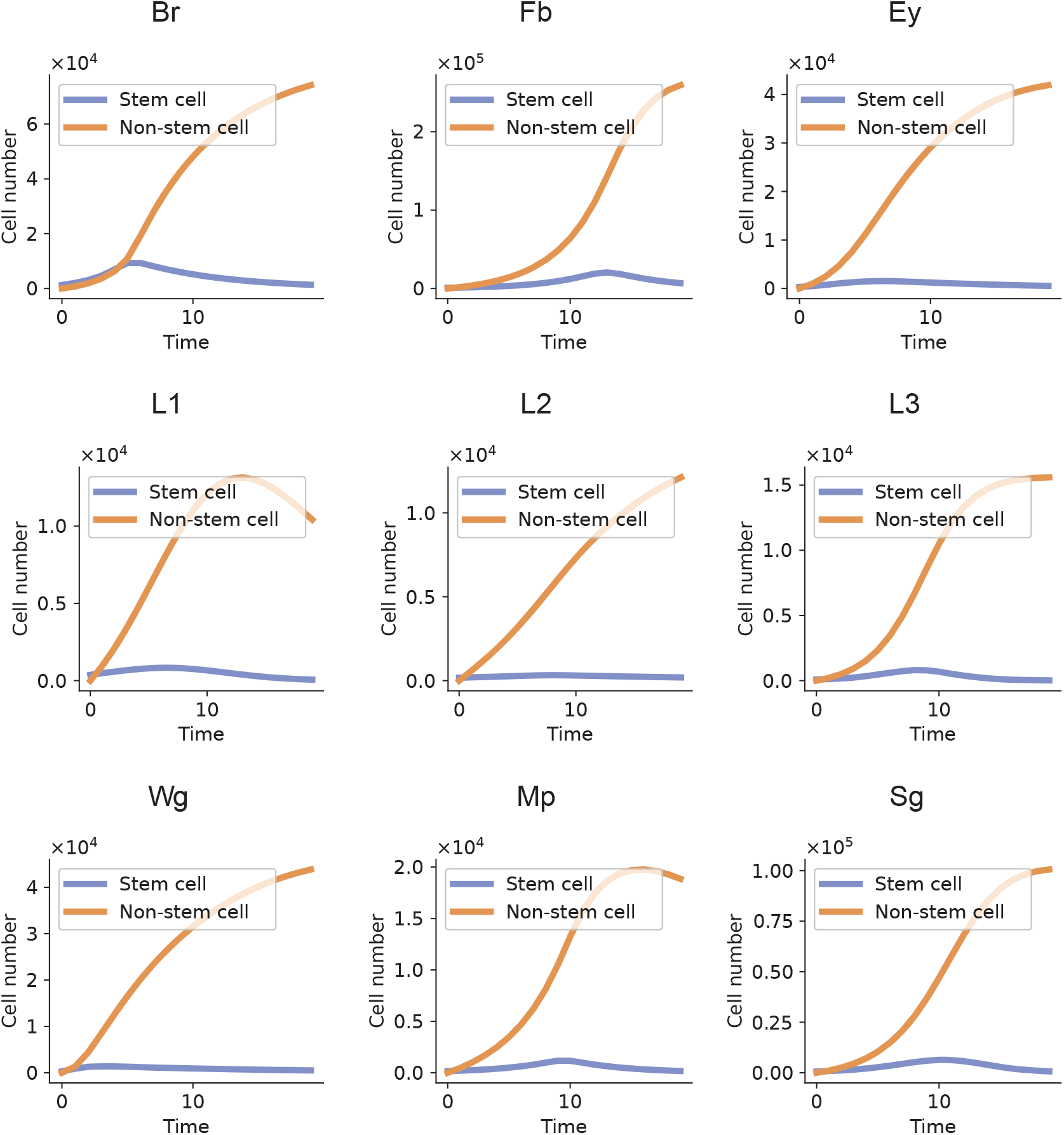
scPhyloX identifies overshoot development in Fly 2. The inferred population growth of stem cells and non-stem cells during development of fly 2. Stem cells and non-stem cells are marked in purple and orange, respectively.

**Supplementary Fig. 13.**
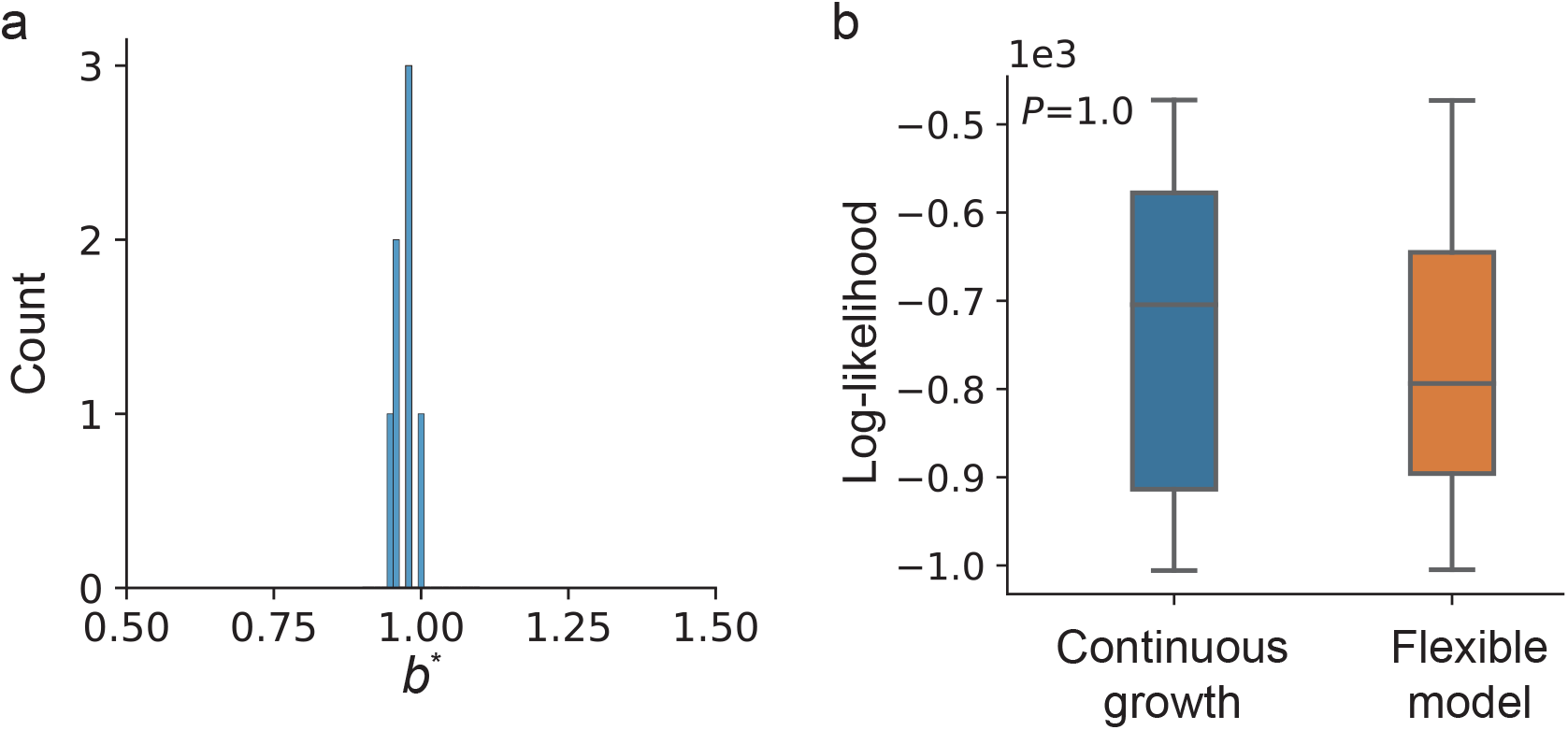
The continuous growth model explains the development of HSC/MPPs better. (**a**) Histogram of inferred *b*^*^ under full parameter model in 8 donors. (**b**) Boxplot shows the model log likelihood between continuous growth model (fixed *b*^*^ = 1) and flexible model. Mann-Whitney U test *P* value shown here.

**Supplementary Fig. 14.**
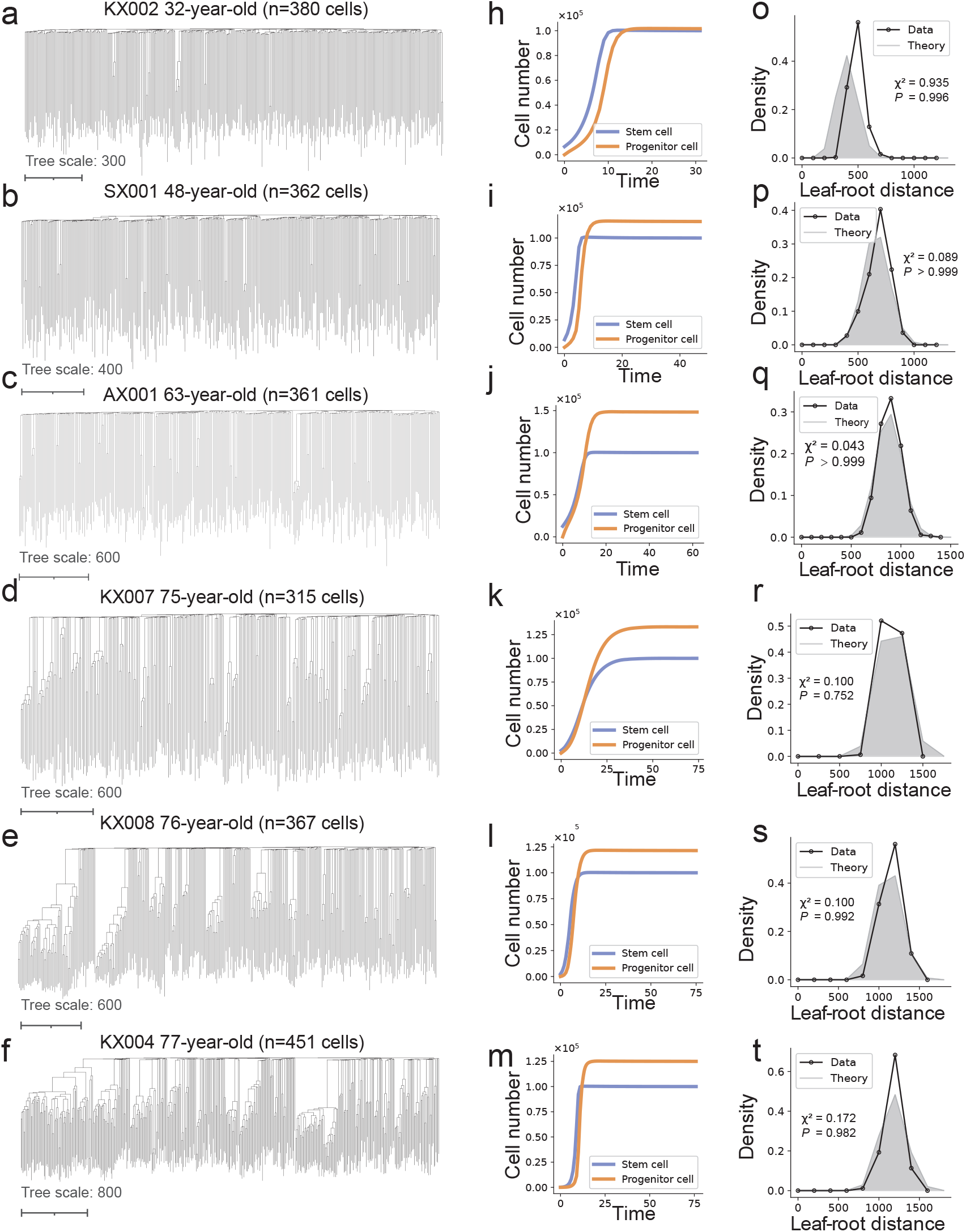
The cell population growth and clonal dynamics during human hematopoiesis. (**a-g**) Phylogenetics tree of HSC/MPPs from donor KX002, SX001, AX001, KX007, KX008 and KX004. (**h-n**) The inferred cell population growth of stem cells (HSC) and progenitors (MPPs) in each sample. (**o-u**) Model fitting of the distribution of leaf-root distances in these individuals, respectively. Pearson’s chi-square tests are shown.

**Supplementary Fig. 15.**
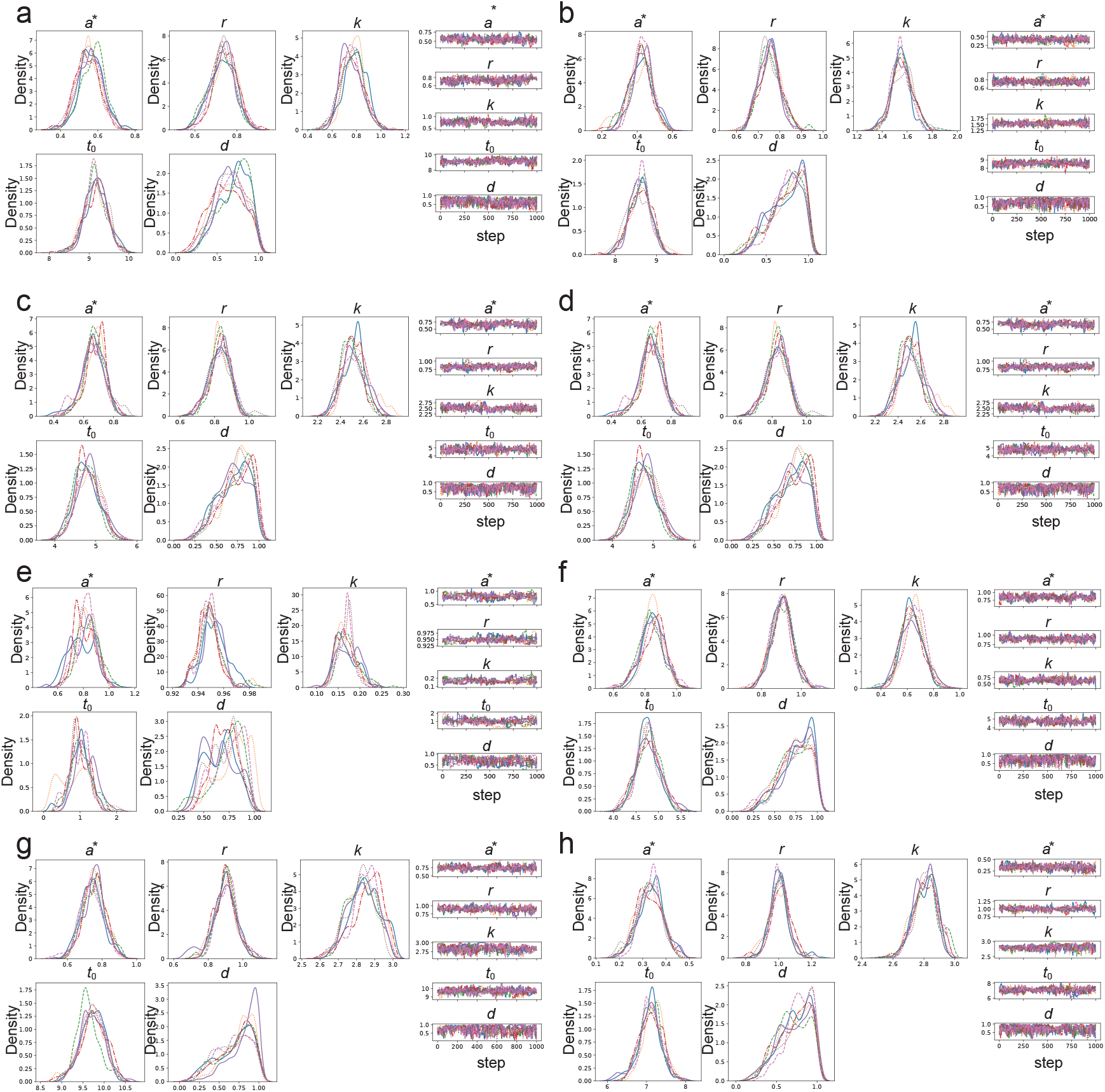
MCMC inference details of HSC/MPPs. (**a-h**) MCMC inferred HSC/MPPs dataset phylodynamics parameter distribution (left) and sampling trace (right) of each donor.

**Supplementary Fig. 16.**
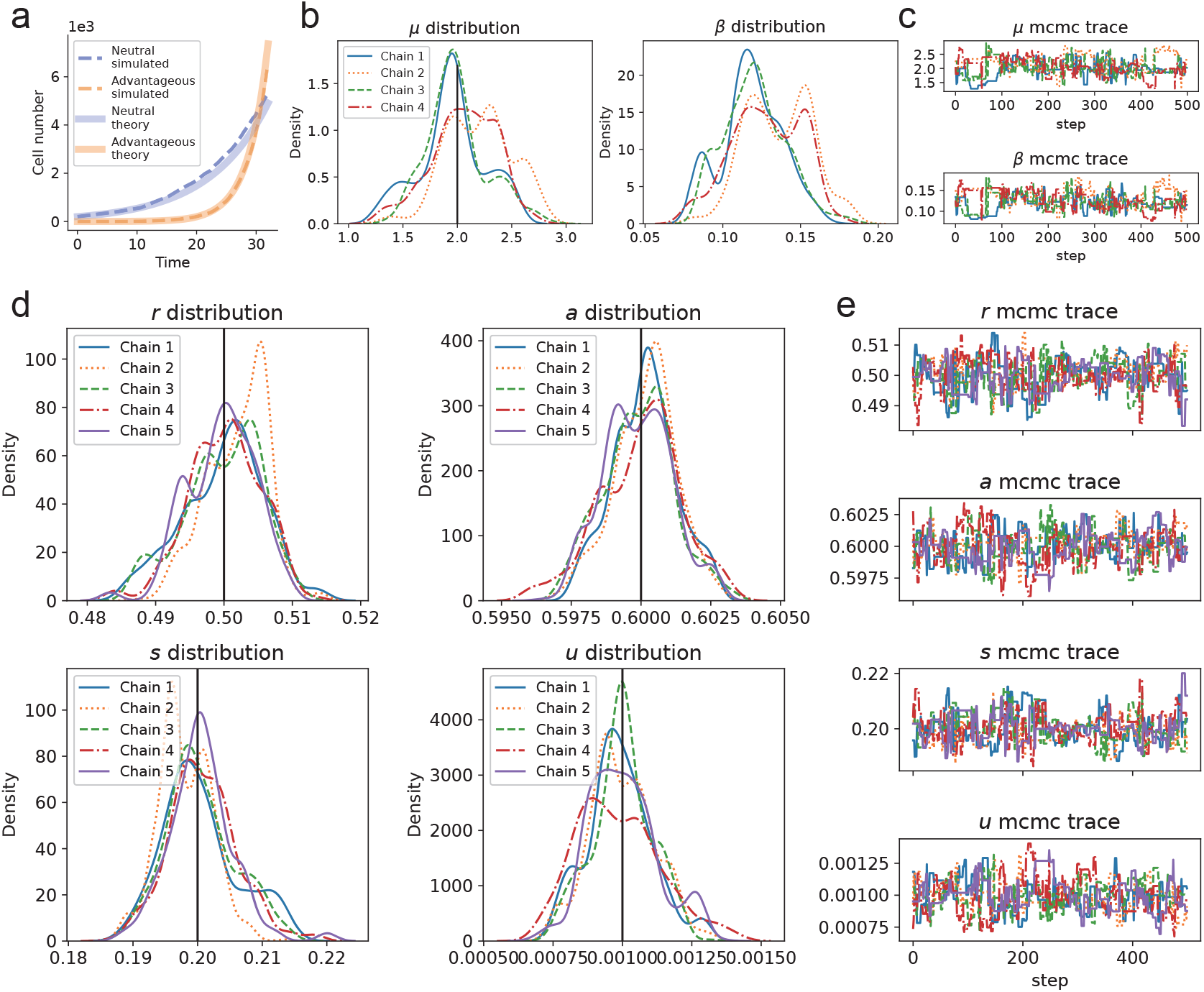
Tumor growth models simulation results and parameter inference details. (**a**) Comparison of the tumor growth model simulation results (dashed line) and the theoretical results (soiled line). (**b**) Posterior distribution of mutation rate estimation. (**c**) MCMC sampling trace of mutation rate estimation. (**d**) Posterior distribution of phylodynamics parameters. (**e**) MCMC trace of phylodynamics parameters.

**Supplementary Fig. 17.**
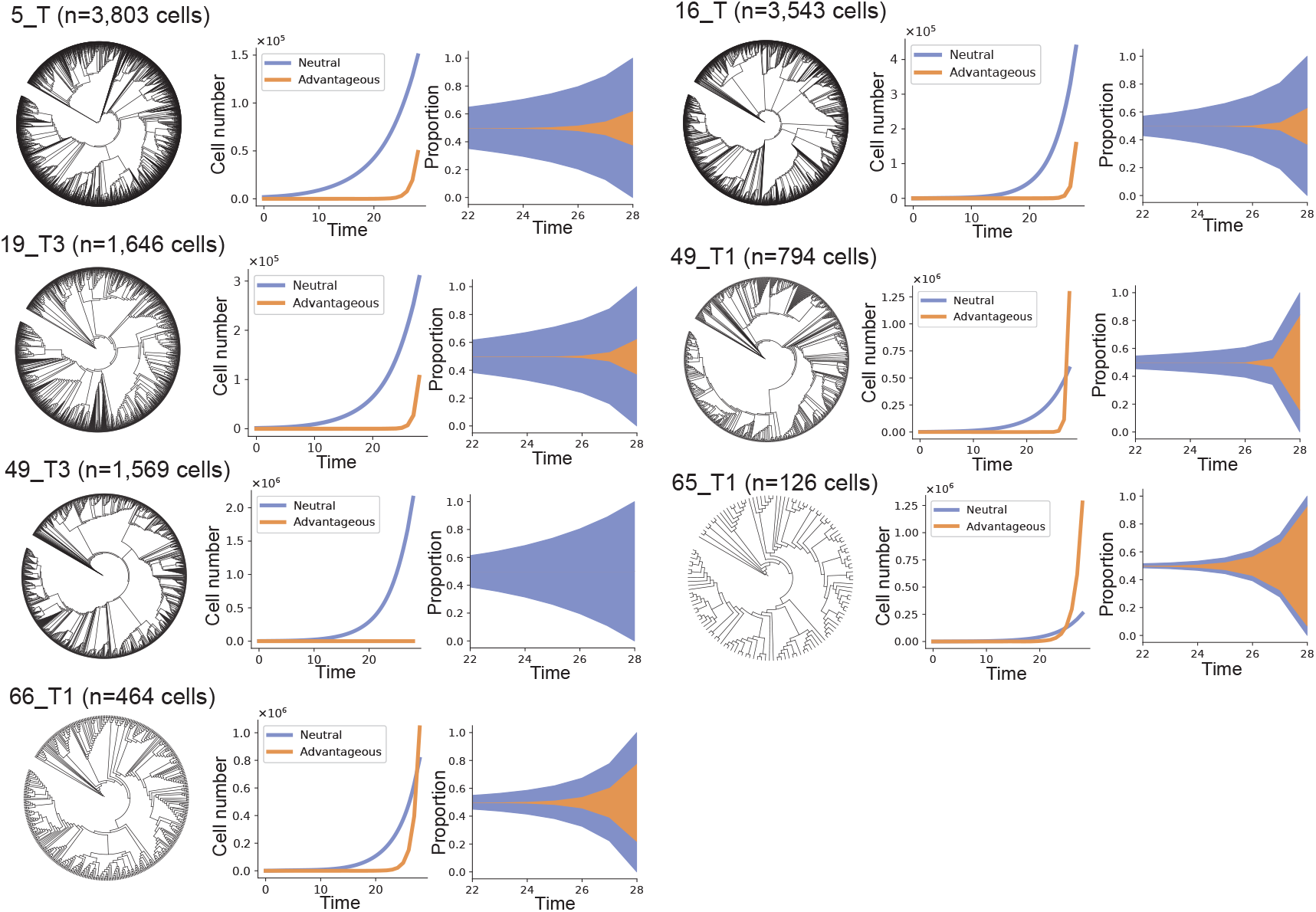
scPhyloX infers tumor growth pattern in early colorectal tumorigenesis. Phylogenetic tree (left), inferred cell population growth of neutral cells (blue) and advantageous cells (orange) (middle) and Muller plot (right) of mouse tumor sample 5 T, 16 T, 19 T3, 49 T1, 49 T3, 65 T1 and 66 T1.

**Supplementary Fig. 18.**
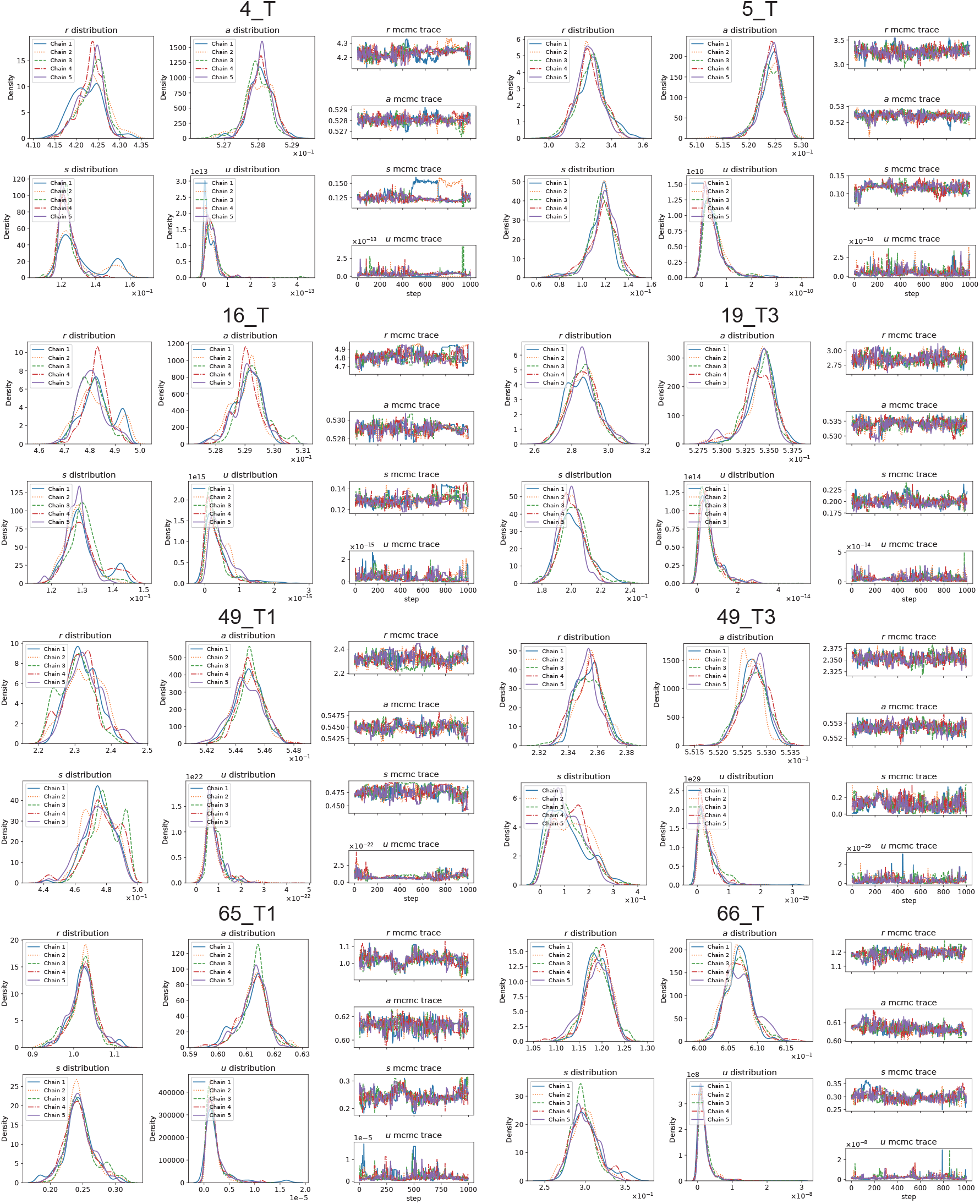
MCMC inference details of tumor growth. MCMC inferred mouse CRC tumor phylodynamics parameter distribution (left) and sampling trace (right) of each sample.

